# Prolonged Ocular Exposure Leads to the Formation of Retinal Lesions in Mice

**DOI:** 10.1101/550236

**Authors:** Brent A. Bell, Vera L. Bonilha, Stephanie A. Hagstrom, Bela Anand-Apte, Joe G. Hollyfield, Ivy S. Samuels

**Affiliations:** Cole Eye Institute/Ophthalmic Research, Cleveland Clinic, Cleveland, OH, United States; Cleveland Clinic Lerner College of Medicine of Case Western Reserve University, Cleveland, OH, United States; Louis Stokes Cleveland VA Medical Center, Cleveland, OH, USA

**Keywords:** mice, retina, lesion, abnormality, imaging, retinal pigmented epithelium, ischemia reperfusion injury

## Abstract

The observation of retinal lesions in the posterior pole of laboratory mice has been found to occur for many reasons, some of which are due to native, developmental abnormalities and those that are influenced by environmental or experimental conditions. Herein, we investigated the rate and extent of retinal lesions as a result of prolonged ocular exposure following general anesthesia. Mice were housed under standard animal care conditions and transported to the laboratory for experimental preparation induction procedures (EPIP) involving general anesthesia, mydriasis/cycloplegia, and topical anesthesia to the cornea. Following EPIP, two ocular recovery conditions (protected and unprotected) were tested within two different animal recovery chambers (open or closed). During anesthesia recovery, and extending up to 2.5 months thereafter, the anterior and posterior poles were evaluated using digital color photography, scanning laser ophthalmoscopy, and spectral-domain optical coherence tomography to document the effects of eye protection and chamber recovery type on the development of retinal lesions. In some mice, electroretinograms and histological evaluations were performed to assess functional and structural changes that accompanied the retinal lesions detected by *in vivo* imaging. We found that the anterior segments of mice recovered in the open chamber with unprotected eyes showed substantial acute changes. At 1-hour post-EPIP, the anterior chamber exhibited corneal thinning, severe media opacities, a reduction in anterior chamber depth, and ocular lens prolapse. These changes largely resolved upon recovery. At 3- and 14-days post-EPIP, inspection of the posterior pole by fundus imaging revealed prominent lesions in the outer retina in a significant proportion of mice recovered in the open chamber. ERG testing conducted at 1-month post-EPIP revealed compromised functional responses in the eyes of affected vs. unaffected mice. Imaging at 14-days post-EPIP revealed that the outer retina lesions in affected mice almost wholly resolve over time to nearly insignificant levels. However, data collected at 80-days post-EPIP demonstrates that some lingering effects persist long-term and appear to be confined to the retinal pigment epithelium. In comparison, mice recovered in the closed chamber with unprotected eyes experienced only mild lens opacities at 1-hr post EPIP that cleared following a full recovery from the effects of sedation. Furthermore, protected eyes of mice recovered in either the open or closed chamber were completely devoid of any anterior or posterior pole complications. In sum, prolonged ocular surface exposure to circulating ambient room air leads to significant anterior and posterior segment ocular complications. We interpret these changes to be caused by dehydration and desiccation of the corneal surface of the eye. The most abundant, semi-reversible complication observed was the development of lesions in the outer retina, which had a 90% probability of occurring after 45 minutes of exposure. The lesions largely absolved short-term but some imaging evidence suggests that they may persist months after their initial appearance.

**Disclosures:** B.A. Bell, none; V.L. Bonilha, none; S.A. Hagstrom, none; B.Anand-Apte, none; 14 J.G. Hollyfield, none; I.S. Samuels, none.

**Grant Information:** Research reported in this publication was supported by the National Eye Institute of the National Institutes of Health under award numbers P30EY025585, R01EY016490, RO1EY026181, RO1EY027083, R01EY014240 and R01EY027750, US Dept. of Veterans Affairs Biomedical Laboratory Research and Development Service VA Merit Award I01BX002754, an unrestricted grant from the Research to Prevent Blindness to the Cleveland Clinic Lerner College of Medicine of Case Western Reserve University, Foundation for Fighting Blindness Research Center Grant, The Wolf Family Foundation, the Llura and Gordon Gund Foundation and the Cleveland Clinic. The content is solely the responsibility of the authors and does not necessarily represent the official views of the National Institutes of Health or the US Dept. of Veterans Affairs.

## Introduction

The laboratory mouse (*Mus musculus*) has been used for over a century in vision research (Pinto and Troy 2008) and is a preferred animal model in biomedical research (Gargiulo, Greco et al. 2012) (Krebs, Collin et al. 2017). During a 10-year tenure of *in vivo* ocular imaging sessions performed by a single operator, instances of “abnormal-looking” retina were observed in mice originating from over 15 principal investigators and collaborating laboratories. Some abnormalities were suspected to be naturally occurring, native problems associated with abnormal eye development (Bell, Kaul et al. 2012), while others were introduced by vivarium lighting conditions (Bell, Kaul et al. 2015). Interestingly, one unique abnormality was observed to occur across multiple strains and/or genotypes, research projects, and mouse treatments. Mice undergoing non-invasive experimental testing procedures involving general anesthesia exhibited similar-looking retinal abnormalities at a relatively low rate of occurrence. These abnormalities were not subtle and could be easily observed with multiple imaging platforms and modalities including color fundus photography, confocal scanning laser ophthalmoscopy, and spectral-domain optical coherence tomography. In some studies, anomalies were observed unilaterally and sometimes bilaterally in about 15-25% of subjects. Examples of procedures where mice developed abnormalities included laser-induced choroidal neovascularization, electroretinography, and various experimental treatments involving the administration of pharmaceuticals via subcutaneous or intraperitoneal injections, herein referred to as “primary” procedures. All occurrences shared a common trend in that they involved: 1) administration of an injectable agent for achieving general anesthesia, 2) administration of topical drops for pupil dilation and topical anesthesia, 3) experimentation involving one of the aforementioned “primary” procedures, and 4) a post-session recovery period.

To determine whether these abnormalities may be pre-existing (Bell, Kaul et al. 2012), baseline imaging was performed in some cohorts of mice prior to the start of the primary experiments. Abnormalities were not observed during this pre-screening session thus ruling out the possibility that these particular ocular complications were the result of preexisting conditions. However, continuation of “primary” procedures using baseline-screened mice would again result in the development and appearance of retinal abnormalities. Following these episodes, it became clear that the procedures and/or conditions that mice were experiencing within the “primary” procedures resulted in the development of the retinal abnormalities.

We initially speculated that the abnormalities originate from pharmaceutical-induced exophthalmia. For decades, mice undergoing laboratory experiments have been administrated the popular drug combination Ketamine and Xylazine (KX) for general anesthesia (Arras, Autenried et al. 2001, Gargiulo, Greco et al. 2012). Xylazine is an alpha-2 adrenergic receptor agonist that has been reported to induce exophthalmia in mice and rats, either systemically via injection, or topically by direct application to the eye (Calderone, Grimes et al. 1986, Zeller, Meier et al. 1998). Mice anesthetized with another less popular small animal anesthesia agent, Sodium Pentobarbital (NaP), do not exhibit a similar proptosis effect. However, we observed that mice administered a very small amount of phenylephrine after NaP develop exophthalmia similar to that of KX anesthetized animals. Phenylephrine, a routinely employed clinical mydriatic, is yet another adrenergic receptor agonist that acts in a dose dependent manner to induce proptosis on the mouse eye after being topically administered to the cornea (see **Suppl. Fig. 1**).

Xylazine and Phenylephrine have been routinely used in combination to prepare mice for ophthalmology-related experiments that require both anesthesia and pupil dilation. Given that these drugs act similarly on the same general receptor and on the eye as a whole by inducing proptosis, we sought to determine the possible consequences of using them in combination. We hypothesized that when used together excessive proptosis occurs secondarily to agonist-induced extraocular muscle relaxation and vasoconstriction, which may initiate retinal abnormalities.

In the course of experiments conducted to test this hypothesis, it became clear that protecting the eye after experimental procedures mitigated ocular complications and that the drug-induced action involving proptosis was not the sole underlying trigger for the development of the observed retinal abnormalities. When performing additional studies to elucidate this phenomenon, we found that retinal abnormalities can frequently occur without careful post-procedural ocular care. The studies presented here will hopefully assist others in ensuring that (1) retinal abnormalities are eliminated from all studies where they are not desired and could ultimately lead to confounding results, and (2) to potentially offer the vision research community an interesting new acute model of localized outer retinal damage that can be non-invasively induced without the need for ocular surgery.

## Methods

### Animal Subjects

Forty-four wild type mice were obtained from the Cole Eye Institute animal vivarium under approved animal use protocols by the Cleveland Clinic Lerner College of Medicine Institutional Animal Care and Use Committee. The experimental procedures described herein were in accordance with the ARVO Statement for the Use of Animals in Ophthalmic and Vision Research. Approximately half the mice were *Tulp1^+/+^* (n=20, 1:1 male/female; Age: 10-39 wks.) on a *C57BL/6J* background (Hagstrom, Duyao et al. 1999) and the other half *C57BL/6J* (n=24, 1:1 male/female; Age: 10-25 wks.) (The Jackson Laboratory, Bar Harbor, ME). Both lines tested negative for the Rd8 mutation of the Crb1 gene (Mattapallil, Wawrousek et al. 2012) as previously described (Bell, Kaul et al. 2015). All mice were housed on ventilated cage racks under standard vivarium conditions including a 14:10 hour cyclic lighting, food and water ad libitum, corncob bedding, and cotton fiber nesting square and red-translucent enrichment hut. Experiments were performed over a 3-month period in 7 groups of mice.

## Experimental Induction of Retinal Lesions

### Procedure 1: Uninterrupted Recovery Experiments

Mice were anesthetized using a mixture of Ketamine (80 mg/kg) and Xylazine (16 mg/kg) diluted in 0.9% saline to replicate routine experimental procedures requiring deep sedation (e.g. surgery, ocular imaging, electroretinograms, drug injections, laser-induced choroidal neovascularization induction, etc.). Within minutes of sedation, mydriasis/cycloplegia and topical anesthesia was induced by administering single drops of 2.5% Phenylephrine (Akorn Inc., Lake Forest, IL, USA), 0.5% Proparacaine, 1% Tropicamide, and 1% Cyclopentolate (Bausch and Lomb, Tampa, FL, USA), applied consecutively to the cornea. The process of inducing general anesthesia, mydriasis/cycloplegia, and topical anesthesia will herein be referred to as an Experimental Preparation Induction Procedure (EPIP). Approximately 1-2 minutes later, right (OD) eyes were protected by receiving a liberal dose of PuraLube Vet Ointment (Dechra Veterinary Products) used in conjunction with an ocular eye shield (Bell, Kaul et al. 2014) whereas left (OS) eyes remained unprotected for the duration of the experiment.

Mice were placed into one of two acrylic containment devices for recovery, herein referred to as the “open” and “closed” chambers. Both chambers (Surgivet V711801, Smiths Medical, Dublin, OH, USA) were placed directly atop a heated hard pad connected to an Androit Medical Heat Therapy Pump (Braintree Scientific, Braintree, MA, USA). The open and closed chambers were exposed or isolated to the ambient room environment, respectively. Temperatures and relative humidity were documented using an indoor/outdoor digital thermometer and digital volt/temperature meter (Extech Instruments, Waltham, MA) with a Type T thermocouple. The open chamber had a temperature range of 21-23ºC and relative humidity of 30-45%. The closed chamber had a temperature range of 28-30ºC and relative humidity of 75%-95%. High relative humidity was maintained in the closed chamber by placing a moist paper towel on the bottom in addition to percolating dry compressed air through a custom-fabricated nebulizer. An appropriately sized silicone finger matt (Ambler Surgical, Exton, PA) was placed on top of moist paper towels to prevent mice from aspirating water condensate. At one hour post-sedation, mouse abdominal surface temperatures were measured (mean±SD) and found to be 32.6±2.9ºC (n=5) and 35.5±0.8ºC (n=7) for the open and closed chambers, respectively. Mice were permitted to recover uninterrupted until regaining consciousness as assessed by evidence of mobility. The approximate time required for recovery was documented individually for each mouse.

### Procedure 2: Interrupted Recovery Experiments

Eight mice that did not develop retinal lesions in the uninterrupted experiments were recycled for use in interrupted recovery experiments. In these experiments, mice were prepared as aforementioned in an identical manner, albeit initially without ocular protection. Immediately post-EPIP mice, mice were placed into the open chamber and permitted to recover naturally from the effects of anesthesia until exposure durations of 25, 45, 65 or 75 minutes were reached. Exposed eyes were subsequently covered with ointment and eye shields upon completion of the exposure duration process. Using a total of 8 mice, four eyes were tested at each exposure interruption time.

### Ocular Imaging

Various imaging modalities were employed to capture the ocular changes occurring to both the anterior and posterior segments. Images were collected immediately following EPIP and for up to 1.5 hrs afterwards during the uninterrupted and interrupted recoveries. Animals were temporarily removed from their respective chambers, imaged and returned as quickly as possible.

Follow-up imaging to assess for the presence or absence of retinal lesions was performed at 3 and 14-days post-EPIP. Images were collected as previously described (Bell, Kaul et al. 2014). A small number of mice were followed for up to 2.5 months to assess whether lesions persist long-term.

### Digital Color Photography

An Apple iPhone 6+ was used to capture the effects of ocular protection or exposure on eyes in both the open and closed recovery chamber conditions. Images were collected from each mouse under standardized conditions that included a front-facing photo collected from a fixed 4” distance with the following settings (50% zoom, HDR On, original color setting, no flash, autofocus frame positioned on mouse forehead). Overhead room lighting was neutral white (4000K) LED room lighting and measured to be ~500 Lux at bench top level.

Examples of retinal lesions at 3 and 80 days post-recovery were also captured using a custom-made topical endoscope fundus imaging (TEFI) system previously described (Paques, Guyomard et al. 2007).

### Confocal Scanning Laser Ophthalmoscope (cSLO or SLO)

A model HRA2 SLO (Heidelberg Engineering, Franklin, MA) was used to collect retinal fundus photos using 6 imaging modes including Infrared reflectance (IR), Infrared Dark-field (IRDF), Infrared autofluorescence (IRAF), Blue autofluorescence (BAF), Red Free Dark-field (RFDF) and Sodium Fluorescein Angiography (FA). A 55° wide-field lens was used to collect images with the optic disk centrally located in addition to peripheral views of the various regional quadrants.

### Spectral-Domain Optical Coherence Tomography (SD-OCT)

Anterior and posterior pole imaging was performed using a Bioptigen model SDOIS SD-OCT system (Leica Microsystems, Buffalo Grove, IL). A Bioptigen mouse bore objective lens with a 50° field of view (FOV) was used for posterior pole imaging with an estimated lateral FOV of ~ 1.5 mm. Imaging of the anterior pole was performed using a 1-inch telecentric lens with an *en face* FOV of 5 mm (azimuth) x 5 mm (elevation) x 3.2 mm (depth). Orthogonal B-scans of the anterior and posterior poles were collected using a radial volume scan (1000 A-scans/B-scan; 2 B-scans, 15 frames). For the anterior pole, scans were positioned just inferior and to the side of the corneal apex reflex to avoid capture of streak artifact from bright specular reflections. Images of the posterior pole were collected at the horizontal and vertical meridians with the optic disk centrally positioned. Additional images of peripheral regions were collected in order to capture retinal pathology examples as needed.

### Electroretinograms (ERG)

Photopic and scoptic electroretinography was performed on *C57BL/6J* mice as previously described (Samuels, Bell et al. 2013). After overnight dark adaptation, mice were anesthetized with Ketamine (80 mg/kg) and Xylazine (16 mg/kg) diluted in 0.9% saline, the cornea was anesthetized with 1% proparacaine hydrochloride, and the pupils were dilated with 1% tropicamide, 2.5% phenylephrine hydrochloride, and 1% cyclopentolate. Mice were placed on a temperature-regulated heating pad throughout each recording session. Responses of the outer retina were recorded using an Espion E3 ColorDome full-field ganzfeld (Diagnosys, Lowell, MA) with Ag-AgCl cornea electrodes referenced to a needle electrode placed in the cheek of the mouse and a ground electrode in the tail. For scotopic ERG, ten steps of blue (445nm) + green (520nm) light flash stimulus [-3.6 to 2.1 log candela (cd)⋅s/m^2^] were presented in the dark in order of increasing flash strength, and the number of successive trials averaged together decreased from 20 for low-level flashes to 2 for the highest flash stimuli. The duration of the interstimulus interval increased from 4 s for low-luminance flashes to 90 s for the highest stimuli. The amplitude of the a-wave was measured 6.0 ms after flash onset from the prestimulus baseline. The amplitude of the b-wave was measured from the a-wave amplitude at 6.0 ms to the peak of the b-wave. Immediately following the dark-adapted strobe-flash stimuli, a steady 20 cd/m^2^ adapting field was presented in the ganzfeld bowl. After 10 min of light adaptation, photopic ERG recordings were obtained from strobe-flash stimuli (-1 to 2 log cd⋅s/m^2^) superimposed on the adapting field. The amplitude of the b-wave was measured from the prestimulus baseline to the positive peak of the waveform. Statistical significance was determined by using a Multiple T-test corrected for multiple comparisons using the Holm-Sidak method with GraphPad Prism 6.0 software.

### Data Processing and Analysis of Imaging Data

Images were exported from their respective imaging platforms to ImageJ 1.47b (Rasband 1997-2012) and Adobe Photoshop CS5 for processing and display. IPhone 6+ and SLO images were exported as JPEG and TIFF, respectively. Anterior and posterior pole SDOCT images were exported as .AVI files, opened in ImageJ, coregistered and averaged using StackReg/TurboReg plug-ins (Thévenaz, Ruttimann et al. 1998). Graphical display of data and statistical analysis was accomplished using GraphPad Prism 6 (Graphpad Software, La Jolla, CA). Unless noted, all data are shown as mean±standard deviation (SD). For all statistical tests, p values and adjusted p values are shown as actual written numerical values or asterisks as follows: ns = *p* > 0.05; \**p* < 0.05; \*\**p* < 0.01; \*\*\**p* < 0.001; \*\*\*\**p* < 0.0001.

iPhone images of ocular media opacities were analyzed using ImageJ by encircling the pupil and obtaining the mean red, green, and blue (RGB) values using the Measure RGB plug-in. Corneal specular reflections from overhead lighting were omitted from the analysis. RGB data (**Fig. 1A-B)** taken during the uninterrupted recovery experiments was converted to grayscale and analyzed to obtain mean opacity magnitude. An Ordinary One-way ANOVA with Sidak’s Multiple Comparisons test was used for determining statistical significance.

**Figure 1.**
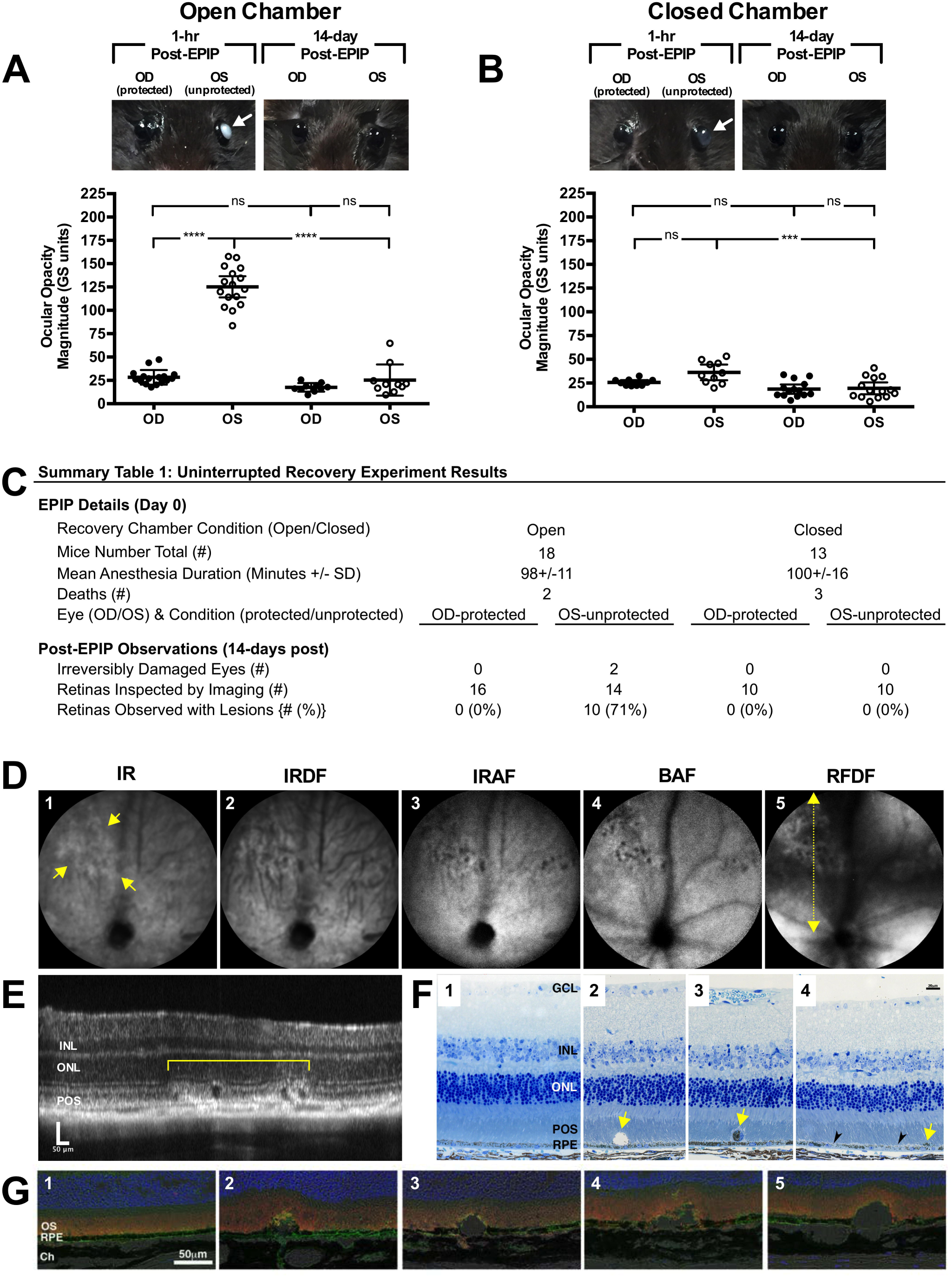
Uninterrupted Recovery Experiment Results. (**A & B**) Mice with eyes that receive no form of ocular protection developed lens media opacities that could be visualized by the naked eye. Opacities were worse in the exposed (OS) eyes of mice recovered in the open (A) vs. closed (B) chamber at 1-hr post-EPIP. For the most part, these opacities resolved by 14-days post-recovery with exception to two instances of irreversibly damaged eyes that resulted in microphthalmia. Eyes that were covered with protective ointment and eye shields did not develop any significant lens media opacities regardless of whether the recovery chamber was open or closed. Arrows indicate the eyes with visible opacities. Note that there is a distinctive difference in the appearance of the two opacities between mice recovered under high-humidity conditions of the closed chamber vs. a typical room environment with low humidity levels present in the open chamber. (**C**) Experimental details and summary table of the observations made for fundus imaging of mice at 14-days post-EPIP. (**D**) Representative SLO images from a mouse with retinal lesions using IR, IRDF, IRAF, BAF, and RFDF imaging modes. Yellow arrows indicate the margin of a retinal lesion observed 14-days post EPIP recovery. RFDF (**D**; yellow dotted-line w/arrows) image indicates the approximate location of the SD-OCT B-scan shown (**E**) that was collected through an SLO detected lesion. Dark spots in BAF-SLO image indicative of the cysts found subsequently by SD-OCT imaging (**E**) and histology (**F^2^ & G^2-5^**). (**F**) Histomicrographs of one normal (**F^1^**) and three abnormal (**F^2-4^**) examples collected from unaffected or affected mice, respectively. **F^2^** shows an enlarged cyst above the RPE that is largely devoid of material with exception to several pigment granules. **F^3^** shows a detached, nucleated cell filled with pigment. **F^4^** shows clustered pigment (yellow arrows) and hypopigmented (black arrowheads) regions of RPE. Immunohistomicrographs of one normal (**G^1^**) and four abnormal (**G^2-5^**) examples collected from unaffected and affected mice, respectively (Blue-TO-PRO-3, Green-GLUT1, & Orange-Rhodopsin). (**G^2-5^**) Sub-retinal cysts and disruptions to the photoreceptor outer segments and interface with the RPE are readily visible. Perturbations as large as 50 μm can be seen displacing photoreceptor inner and outer segment lamina primarily in the vitreal direction.

SLO images of the retina were analyzed for average lesion count or number (#), individual lesion size (area%), and collective or total, accumulative lesion area (Σarea%) for the available image FOV. The available FOV for the uninterrupted recovery experiments included both central and peripheral SLO 55° views that included the superior, temporal, and nasal retinal regions, all of which were analyzed independently. Central view data (**Suppl. Fig 2B**) was analyzed with an unpaired two-tailed t-test with equal standard deviations. Peripheral view data (**Suppl. Fig 2C**) was analyzed with an ordinary One-way ANOVA with Tukey’s multiple comparisons test.

The available image FOV for the interrupted recovery experiments was a Photoshop CS5 montage that combined central and peripheral views collected with the SLO 55° wide-field lens from all four retinal quadrants into a single image of the retina. Lesion count (#), individual lesion size (area%), and collective or total, accumulative lesion area (Σarea%) (**Fig. 2A**) obtained from the montaged images were analyzed using an ordinary One-way ANOVA with Tukey’s multiple comparisons test.

**Figure 2.**
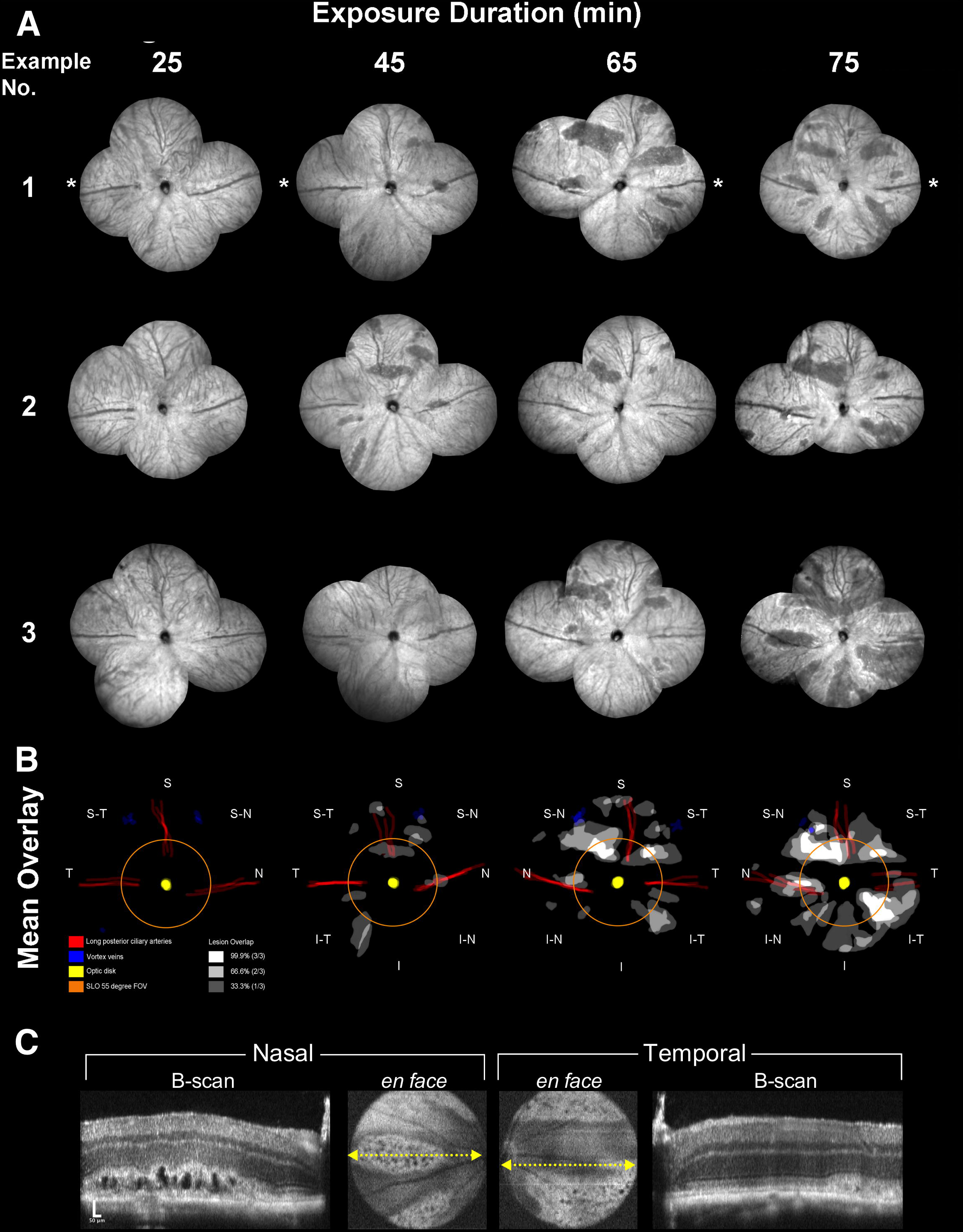
Montaged fundus views and results from the interrupted exposure time point experiments. The extent of the induced abnormalities at 3 days post-EPIP can be seen in three examples provided for each interruption time point (**A**). Asterisks indicate the temporal region. No abnormalities are visible at 25 minutes whereas the number and size of the lesions can be seen increasing with longer exposure duration. Lesion involvement overlays were created for each retina example that also included demarcating the location of the vortex veins, optic disk, and long posterior and superior ciliary arteries. Mean overlays of lesion involvement were created to identify lesion development hot spots related to retinal quadrant (**B**). As observed, lesions occur in various regions of the posterior pole and in particular, within specific lanes or zones within those regions. The orange ring shows the extent of FOV for the SLO 55° wide-field lens. Note that many lesions form outside this central retina FOV, which is fairly conserved among commercially available color fundus, SLO and SD-OCT imaging instruments capable of imaging mice. Based on the visible appearance of the lesions within these montages it is suspected that they extend further out beyond the FOV and presumably impacting retina possibly out to the ora serrata in some instances. (**C**) SD-OCT images from the horizontal meridian of the mouse from **Fig. 2A** IRDF-SLO example #3 @ 75 minutes Post-EPIP recovery time showing prominent formation of cysts within the photoreceptor layer.

SLO fundus image overlays were compiled using montaged images from the interrupted recovery experiments to better identify regions where lesions had the highest tendency to materialize. Using Photoshop, features of the retina were filled or traced, including the lesion involvement area (white), long-posterior ciliary arteries (red), optic disk (yellow), and vortex veins (blue) **(Fig. 2B)**. Montaged images from three mice at exposure durations of 25, 45, 65, 75 minutes were overlaid and aligned in Photoshop using the long-posterior arteries and optic disks as landmarks. Once combined, the three individual montages from each time point were averaged to obtain a heat map of area overlap **(Fig. 2B)** showing the areas of highest (white) and lowest (black) tendencies for lesion formation.

Anterior segment SD-OCT data was analyzed for exophthalmia (e.g. proptosis), cornea thickness, anterior chamber depth, lens media opacity area, lens media opacity magnitude, and lens media opacity integrated density using ImageJ. Exophthalmia was measured from the cornea apex to the medial canthus. Cornea thickness and anterior chamber depth was measured from horizontal and vertical orthogonal B-scans through the central optical axis and averaged over ~10 frames. Lens media opacity data was obtained by encircling the opacity using a drawing tool and obtaining area, magnitude, and integrated density values in ImageJ. Area measurements of media opacities were converted from pixels to square millimeters by using a ruler for calibration of the B-scan image frame. Scatter plots were generated using pooled data from both uninterrupted and interrupted recovery experiments and non-linear regression curve fits were performed to show mean±95% confidence interval bands. Data were fitted to a best-fit curve using R-squared values, which was usually a straight line or one-phase exponential decay (or association).

RGB values collected from the media opacity images (**Fig 7A)** obtained during the interrupted recovery experiments were converted to CIE 1976 L*a*b* (Lab) color space (**Fig. 7B-D**) using an online conversion tool (Colormine.org). Lab hexadecimal color values were obtained using an online color picker tool (DavidJohnstone.net). MS Powerpoint was used to generate color bars and mouse pupil color replications using the obtained color values. Statistical significance for the mean Lab values measured was determined using an ordinary One-way ANOVA with Tukey’s multiple comparisons test.

A Pearson correlation test was performed only for mice/eyes that developed retinal lesions. The test obtained correlation coefficients and p-values between collective lesion impact area and other measured variables that included a total of 236 total data points for exposure duration, exophthalmia, cornea thickness, anterior segment depth, lens media opacity area, magnitude and integrated density, and CIE L*a*b* values.

### Histology

Eyes were enucleated and fixed by immersion in 2% paraformaldehyde, 2.5% glutaraldehyde and 5% CaCl_2_ made in 0.1 M cacodylate buffer overnight at 4°C and processed for epon embedding and imaging as previously described (Bonilha, Bell et al. 2015). Semi-thin sections were cut with a diamond histotech knife, collected on glass slides, and stained with toluidine blue. Slides were photographed with a Zeiss AxioImager.Z1 light microscope and AxioCam MRc5 camera.

### Immunocytochemistry

Eyes were enucleated and fixed by immersion in 4% paraformaldehyde in PBS at 4°C, quenched with 50 mM NH_4_Cl for 30 min and then infused successively with 10% and 20% sucrose in PBS, and finally Tissue-Tek “4583” (Miles Inc., Elkhart, IN). Cryosections (8 μm) were cut on a cryostat HM 505E (Microm, Walldorf, Germany) equipped with a CryoJane Tape-Transfer system (Leica Inc., Buffalo Grove, IL). For labeling, sections were washed to remove embedding medium, blocked in PBS supplemented with 1% BSA (PBS/BSA) for 30 min, and incubated with primary followed by secondary antibodies coupled to Alexa 488 or Alexa 595 and finally incubated with TO-PRO-3 for nuclear labeling (LifeTechnologies, Grand Island, NY) as previously described (Bonilha, Bell et al. 2015). A series of 0.3-μm *xy* (*en face*) sections were collected using a laser scanning confocal microscope (Leica TCS-SP8, Exton, PA) using the same acquisition parameters for each channel in the Leica confocal software LAS AF. Antibodies used included rhodopsin (clone B630N, from Dr. G. Adamus, Oregon Health and Science University, Portland, OR, 1:100), glucose transporter GLUT1 antibody (ab652, 1:200).

## Results

### Uninterrupted Recovery Experiment Results

**Figure 1** shows the observations and data obtained from the Uninterrupted Recovery Experiments. Right eyes (OD) that were protected were noticeably absent of any visible evidence of media opacity regardless of the recovery chamber condition. In contrast, left eyes (OS) that experienced prolonged ocular exposure exhibited both visible and quantifiable differences in media opacity response. As discerned in the digital photographs (**Fig. 1A & 1B**; white arrows), mice recovering in the open (126.9±21.9 grayscale or “GS” units) vs. closed (36.2±11.5 GS units) chambers with unprotected eyes were significantly (P<0.0001; unpaired two-tailed t-test) more prone to developing severe media opacities. All ocular media opacities resolved to insignificant levels following recovery (**Fig. 1A & 1B**) when the mice were re-evaluated for evidence of retinal lesions at 14 days post-EPIP. However, two mice from the open chamber with unprotected left eyes developed corneal ulcerations and microphthalmia that prevented retinal imaging assessment.

In two cohorts of wild-type mice tested, posterior pole imaging at 14-days post-EPIP revealed retinopathy-like lesions in 70% (7/10) and 75% (3/4) of *Tulp1^+/+^* and *C57BL/6J* mice, respectively. Collectively, 71% (10/14) of wild-type mice developed lesions after being under general anesthesia for 1.5 hrs (**Figure 1C)**. A comprehensive analysis of the SLO imaging data was performed for the Uninterrupted Recovery Experiments and provided in **Suppl. Fig. 2A-C**. Only mice that did not receive ocular protection and recovered in the open chamber had lesions visible by imaging. Lesions could be observed to varying degree using the five native reflectance and autofluorescence SLO imaging modes (**Fig. 1D^1-5^**). Two mice, out of 10 affected, had visible lesions only after the camera head was panned to the peripheral retina (see **Suppl. Fig. 2B**). SD-OCT imaging immediately following SLO revealed lesions confined to the outer retina and almost exclusively to the photoreceptor layer (**Fig. 1E**-bracket). Hypo- and hyper-reflective features visualized by SD-OCT within the SLO identified lesion boundaries resembled pathologies common to models of Age Related Macular Degeneration such as Outer Retinal Tubulation (ORT) (Zweifel, Engelbert et al. 2009) and Reticulated Pseudodrusen (RPD) (Khan, Mahroo et al. 2016). Histology confirmed the presence of outer retinal pathology (**Fig. 1F^2-4^ & 1G^2-5^**) in affected areas confined to the photoreceptor outer segments and RPE. Hyporeflective circular features observed by SD-OCT imaging were found to be atypical of ORTs and more similar to pseudo or subretinal cysts as no cells were encircling the vesicular void (**Fig. 1F^2^ & 1G^2-5^**). Additional observations include the evidence of displaced melanosomes and/or melanin pigment granules (**Fig. 1F^2&4^** – yellow arrows), a detached RPE cell or infiltrating sub-retinal inflammatory cell (**Fig. 1F^3^** – yellow arrows), and RPE hypopigmentation (**Fig. 1F^4^**-black arrowheads).

Electroretinograms performed on the *C57BL/6J* cohort one-month post-EPIP demonstrated that functional changes were correlated with the mice recovered without ocular protection in open chambers (**Suppl. Fig. S3**). In **Suppl. Fig. S3A**, the a-wave amplitude of mice without protection in open chambers is significantly smaller than those recovered with ocular protection. Similarly, the b-wave is significantly smaller in response to high flash stimuli **(Suppl. Fig. S3C).** There is also a trend toward smaller light-adapted responses in mice recovered without protection in open chambers as compared to mice with ocular protection **(Supp Fig S3E).** In contrast, mice that underwent unprotected recovery, but in closed chambers, did not display significant reductions in a- and b-wave amplitudes **(Suppl. Fig. S3B & S3D)** or in light adapted response **(Suppl. Fig. S3F)**.

### Interrupted Recovery Experiment Results

Two groups of *Tulp1^+/+^* mice subsequently underwent a second episode of EPIP, with interrupted ocular recovery occurring at 25, 45, 65, or 75 minutes. All mice were evaluated for retinal lesions three days post-EPIP. IRDF-SLO images from the retinas of affected and unaffected mice are shown in **Figure 2A.** Montaged views of central and peripheral retina are shown with an approximate 110° FOV (55° x 2) taken from the horizontal and vertical meridians. At 3-days post-EPIP, lesions appear as dark areas relative to normal background by IRDF-SLO imaging. Qualitatively it can be observed that the lesion number and area expand with increasing exposure duration. At 25 minutes mice had not developed any retinal lesions. After 45 and 65 minutes, 75% (3/4) of mice developed retinal lesions. At 75 minutes, 100% of the mice that could be imaged (3/3) had retinal lesions while the remaining mouse that could not be assessed had irreversible ocular damage in the form of microphthalmia secondary to an ulcerated cornea.

Mean overlays pinpoint areas of the retina that were vulnerable to lesion development (**Fig. 2B**). After 45 minutes of exposure, it is apparent that lesions are forming in the superior-nasal and inferior-temporal regions. At 65 minutes, lesions in the superior-nasal and inferior-temporal regions expand in coverage with an additional dominant location in the superior-temporal region emerging. Additional, smaller lesions appear in the nasal and superior regions at 65 minutes and by 75 minutes, all observances increased in frequency, magnitude and area, and have become widespread throughout the FOV.

Horizontal meridian SD-OCT B-scans from the same IRDF-SLO fundus image shown in **Figure 2A** (Example No. 3 @ 75 minutes) are shown in **Figure 2C**. A nasal region B-scan shows abnormal outer retina morphology through the middle of the lesion. Lesion severity was more pronounced at 3 days post-EPIP than 14 days (**Fig 2C vs. 1E**). Both hyper- and hypo-reflective changes (**Fig 2C**-nasal B-scan) appeared as hard or soft retinal exudates above subretinal pseudocysts or pyramidal “ghost” pseudodrusen, respectively (Khan, Mahroo et al. 2016). The temporal B-scan is taken at the edge of a lesion and thus absent of the cysts but shows a photoreceptor layer absent of normal architecture and axial displacement of external limiting membrane and IS-OS/ellipsoid zone.

**Figure 3A** graphically illustrates the quantified imaging data obtained from the interrupted recovery experiments. The total lesion area (Σarea%) is shown plotted and fitted with an exponential growth curve (Adj. R^2^ = 0.98) in **Fig. 3A** (insert). Mean total lesion area involvement consistently increased in relation to exposure time. Between 45 and 65 minutes, and 65 and 75 minutes, the total lesion area increased ~2.5 times. A One-way ANOVA indicated the increasing trend in total lesion area was significant (p=0.01) as well as the changes observed between 25 and 75 minutes (p=0.008) and 45 and 75 minutes (p=0.034). The mean number (#) of lesions increased with exposure time and was 0±0, 4.3±3.1, 6.3±1.5, 8±1 for 25, 45, 65, and 75 minutes, respectively. Average individual lesion size (area%), relative to the percentage of the montaged FOV, was 0±0, 0.9±0.6, 1.8±0.9, 3.9±2.6 for 25, 45, 65, and 75 minutes, respectively. For reference, the average size of the optic disk for the twelve fundus montages shown in **Fig. 2A** is ~0.24 % of the montaged SLO FOV. Thus, the mean individual retinal lesion sizes calculated were on average 3.75, 7.5 and 16.3 times larger than the optic disk for 45, 65, and 75 minute exposure times when observed by IRDF-SLO imaging at 3-days post-EPIP.

**Figure 3.**
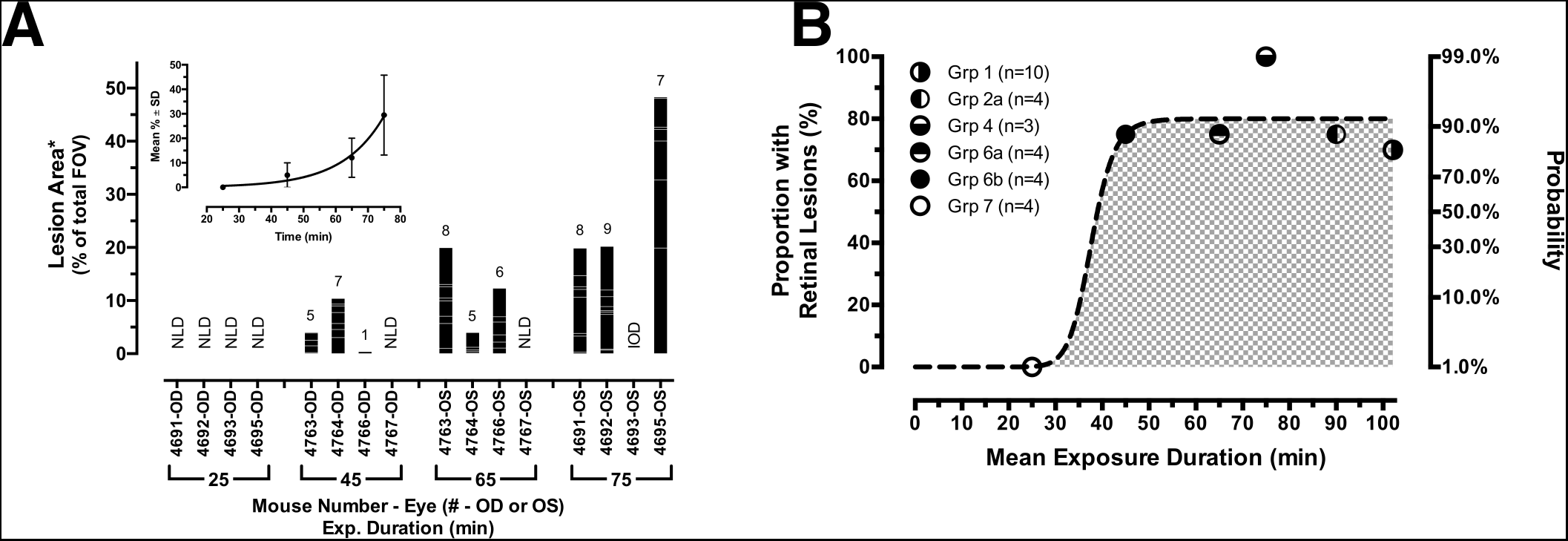
Quantified Results from the Interrupted Experiments. (**A**) Statistics obtained for each mouse eye that underwent interrupted recovery protection at 25, 45, 65, and 75 minutes post-EPIP. All parameters measured from the images shown in **Figure 2** increased with elapsed time and included: (1) the number of animals with lesions, (2) the number of lesions counted per eye, (3) average lesion area and (4) the mean total lesion area (inserted graph). Abbreviations: NLD-no lesions detected, IOD-irreversible ocular damage. (**B**) Exposure time-response curve for all combined mouse data from the uninterrupted and interrupted experiments in eyes that were unprotected when recovered in the open chamber environment. This trend underscores that eyes cannot be left unprotected for more than ½ hour before the risk of developing retinal lesions. Note that the curve is steep and shows that the mice essentially transition from low to high risk within 15 minutes beyond the ½ hour exposure mark. Beyond 45-50 minutes of unprotected exposure, mice have a 90% probability of acquiring retinal lesions.

### Pooled Results Showing Probability of Developing Retinal Lesions

When we combined the results from both the uninterrupted (Grps 1-2) and interrupted (Grps 4,6-7) experiments, a time–response curve could be generated (**Fig. 3B**) for mice with eyes that were unprotected and recovered in the open chamber, that subsequently developed retinal lesions post-EPIP. The primary abscissa shown on the left in the graph corresponds to the proportion of mice per group found with retinal lesions relative to mean exposure duration time for each of those groups. The secondary abscissa shown on the right shows the probability of lesion development relative to exposure time after fitting the data with a sigmoid curve (Adj. R^2^ = 0.84). The effective time estimated for half of the animals to develop lesions (ET_50_) was 37.7 minutes.

### Extended Follow-up of Exposure-induced Retinal Lesions

One mouse observed with lesions at 3-days post-EPIP following a >60 minute recovery was monitored for up to 2.5 months to determine if the acutely induced developments would absolve or persist. **Figure 4** shows examples of lesions documented at 3-days post-EPIP and up to 80 days thereafter by SLO, TEFI and FA-SLO imaging. **Fig. 4A** demonstrates a prominent lesion observed by IRDF-SLO at 3-days post-EPIP that quickly resolves to nearly undetectable levels at 14-days post. Different changes persist at 14-days post, which appear as relatively small perturbations of hypo- or hyper-reflective spots, or variations in background intensity within the original lesion boundary relative to other unaffected areas of the image FOV. The spots slowly absolve over time to non-detectable levels by 80 days post when observed by IRDF-SLO imaging. The same lesion identified by IRDF (**Fig. 4A**) is shown by IR-, RFDF- and BAF-SLO in **Fig. 4B.** The IR reflectance image shows the lesion as a hypo-reflective area in the superior-temporal region that is not readily discernable 80 days post in the superior view; however to an experienced eye, some residual perturbations can be observed, such as punctate hyper- and hypo-reflective spots within the original lesion boundary. In contrast to the two IR imaging modes, blue light illumination imaging modes (RFDF and BAF) showed more apparent features at 80 days post. The RFDF and BAF imaging example suggests an outer retina still actively undergoing modification or repair from the original insult.

**Figure 4.**
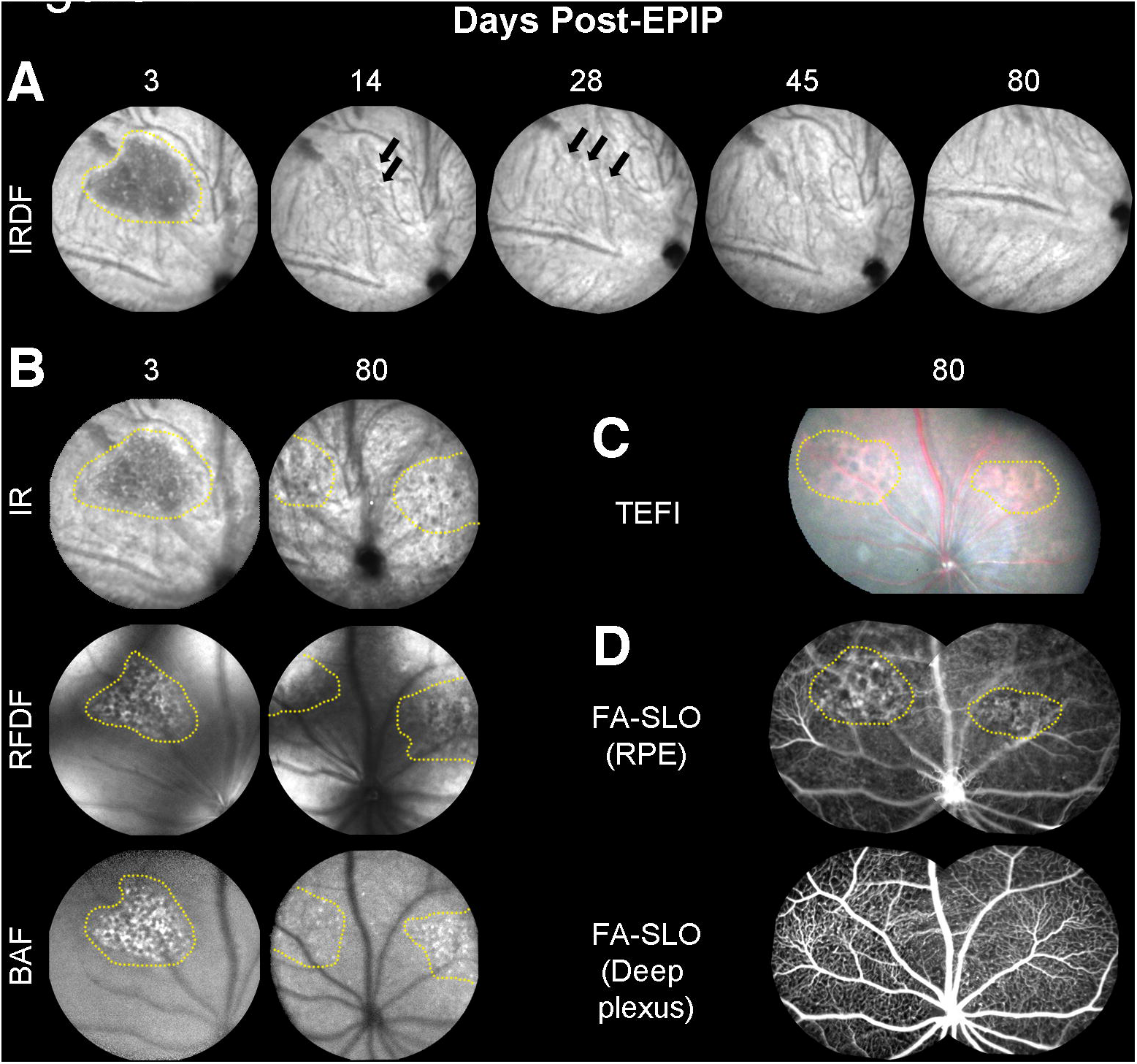
Long-term follow up of retinal lesions. (**A**) IRDF-SLO images of lesions at 3-days post-EPIP, which are easily discernable from the normal RPE/choroidal background. The lesion is presented as a dark region that has resolved 14-days post-EPIP. At 14 and 28-days, subtle indicators (hyper- and hypo-reflective spots) persist that suggest some evidence of the lesion still remains. By 45 and 80-days, the hyper- and hypo-reflective perturbations have resolved and the region appears similar to the surrounding background. (**B**) Additional examples of the same lesion shown by IRDF (**Fig. 4A**) at 3-days post are also readily visible by three of the other SLO imaging modes. (IR, RFDF and BAF). This comparison helps to identify and isolate the region impacted by the lesion, which appears to be the outer retina and RPE. As a result of this observation it is suggestive that the lesion remaining at 80 days post-EPIP is also altered RPE and this is further supported by the TEFI and FA-SLO images that follow. (**C**) Color fundus images showing a different spectral reflectance profile for the two visible lesions versus the surrounding background. (**D**) Sodium Fluorescein Angiography (FA-SLO) revealing leakage and/or abnormal uptake of the fluorophore at the RPE level whereas no evidence of leakage can be observed in the deep plexus of the retinal vasculature. The enhanced visualization of red reflectance and green fluorescence for the TEFI and FA-SLO images, respectively, could alternatively be due to hypo-pigmentation of the RPE as previously indicated in Figure 1F^4^. These observations suggest that the damage remaining at this late stage is perhaps isolated to the RPE.

The same mouse underwent TEFI imaging and SLO angiography to show the appearance of the lesions using visible-light fundus photography and for the presence of sodium fluorescein leakage at the previously documented lesions sites (**Fig. 4C & 4D**). TEFI showed the lesion areas as red in color suggesting visualization of the underlying choriocapillaris and circulating erythrocytes (**Fig. 4C)**. Additional TEFI images of lesions a few days post-EPIP are provided as supplemental materials for comparison to the mature lesion shown (**Fig. 4C**) and demonstrate that recently induced lesions have a reflective white appearance (**Suppl. Fig S4**). FA-SLO of the camera focus trained on the RPE show irregular fluorescence patterns in the super-nasal and super-temporal regions relative to the surrounding areas (**Fig. 4D**). These regions of atypical visualization correspond well to the retinal lesions detected using TEFI and native reflectance/autofluorescence SLO imaging modes that could indicate fluorescein uptake or leakage by the RPE or alternatively, trans-RPE visualization of the circulating fluorescein within the choriocapillaris. Sodium fluorescein demarcation is no longer evident when the camera focus is repositioned to image the deep vascular capillary plexus of the retina further indicating that the defect is isolated to the distal region of the outer retina and perhaps exclusively to the RPE.

### Anterior Segment Dynamics Following EPIP

Imaging data collected as mice were recovering from the acute effects of sedation were also analyzed to investigate what changes occur to the various entities within the anterior segment. **Figure 5** shows the anterior segment changes observed by SD-OCT in mice recovered in open (**Fig. 5A**) and closed (**Fig. 5B**) chambers with eyes that were protected or left unprotected during the recovery period. Similar to the observations made in **Fig. 1A**, eyes that were protected (OD-open & closed chambers) showed very little change compared to eyes that were left unprotected (OS-open & closed chambers). Unprotected eyes in either recovery chamber exhibited media opacities that persisted throughout the duration of the exposure time regardless of chamber recovery type. However, mice recovered in the open chamber had more severe changes to the anterior segment region than those being recovered in the closed chamber.

**Figure 5.**
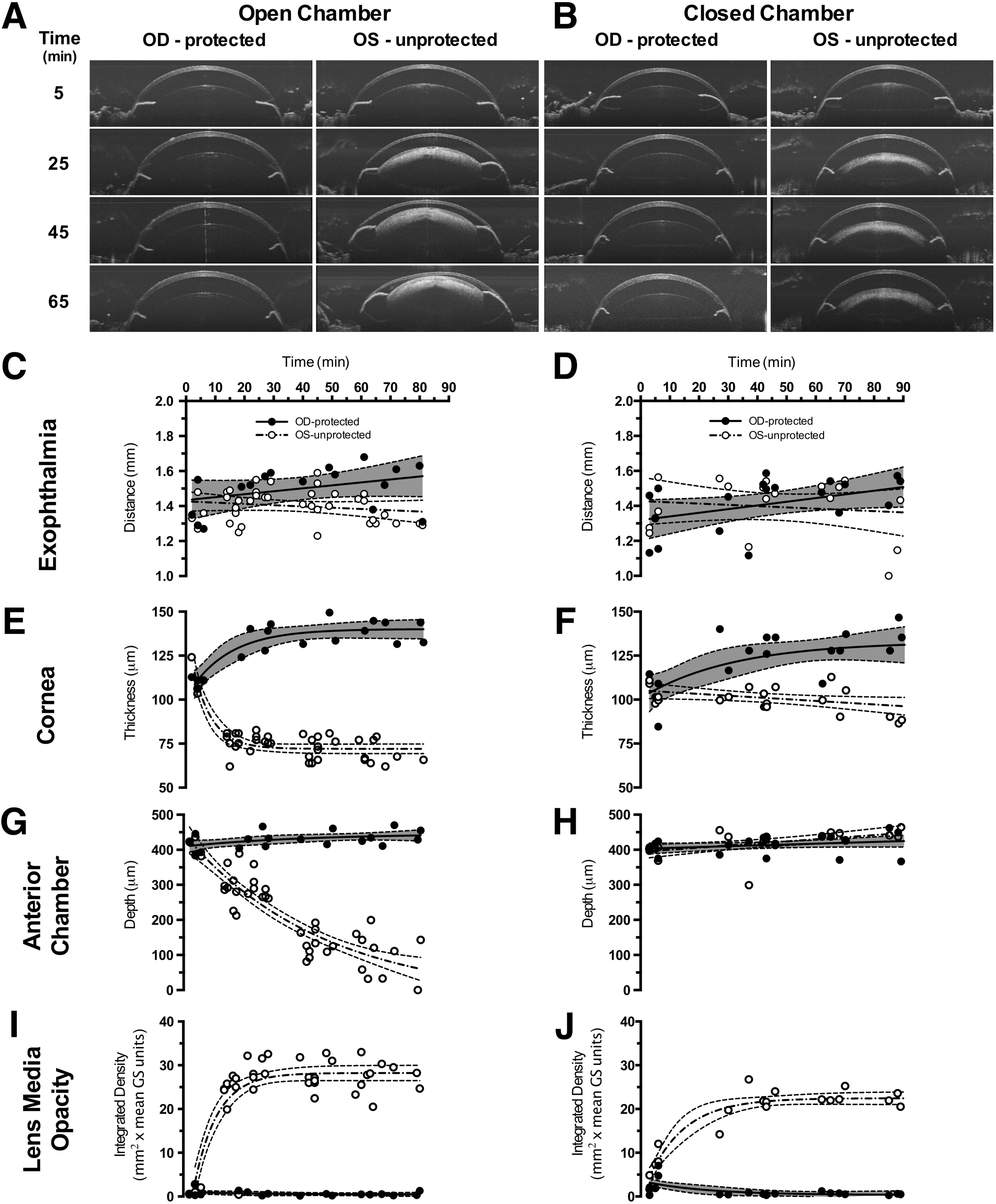
Anterior Segment SDOCT Imaging Examples and Quantified Results. Representative SDOCT images from the anterior segment of mice recovered in the open (**A**) vs. closed (**B**) chambers. Eyes of mice that received ocular protection **(OD-protected)** exhibited no substantial adverse changes compared to eyes that were left unprotected **(OS-protected)**. However, mice with unprotected eyes **(OS-unprotected)** and recovered in the closed chamber (**B**) only developed lens media opacities in contrast to mice with unprotected eyes **(OS-unprotected)** and recovered in the open chamber that (**A**) developed visibly apparent corneal thinning, lens media opacities and reduction in anterior segment depth. Quantitative results for exophthalmia (**C-D**), corneal thickness (**E-F**), anterior chamber depth (**G-H**) and lens media opacity (**I-J**) for the two ocular status and two recovery and conditions.

**Figure 5C & 5D** shows the exophthalmia results which consistently increased (~7-10%) or decreased (~3%) for protected vs. unprotected eyes, respectively, regardless of recovery chamber condition. In the cornea (**Fig. 5E & 5F**), protected eyes showed moderate swelling as thickness increased by 15-20% regardless of recovery chamber condition. Unprotected eyes showed differences in corneal shrinkage trends between mice recovered in closed vs. open chambers. Mice recovered in the closed chamber (**Fig. 5F**) with unprotected eyes exhibited nominal corneal thinning (~7%) whereas mice recovered in the open chamber (**Fig. 5E**) showed substantially more by comparison (~33%). Moreover, mice in the open chamber (**Fig. 5E**) reached this level of change after only 20-25 minutes, which remained an asymptotic limit throughout the remainder of the recovery period. Changes in anterior chamber depth (**Fig. 5G& 5H**) were substantial for one condition, which was for mice recovered in the open chamber without ocular protection (**Fig. 5G,** OS-unprotected). Over the entire recovery period, this group of mice exhibited 400% reduction in anterior chamber depth. In comparison, anterior chamber depths for the other three recovery conditions (**Fig. 5G**, OD-protected and **Fig. 5H**, OD-protected & OS-unprotected) showed nominal increases of only ~10%. No appreciable lens media opacities were identified (**Fig. 5A & 5B; Fig. 5I & 5J**), for protected eyes of mice recovered in either the open or closed chambers. Lens media opacities of unprotected eyes showed qualitative differences between mice recovered in open vs. closed chambers (**Fig. 5A & 5B)**. Integrated density measurements of the media opacities showed that mice with unprotected eyes recovered in the open chamber developed more severe cataracts than mice with unprotected eyes recovered in the closed chamber by about ~20% (**Fig. 5I & 5J)**. **Supp. Fig. S5** separates the products of integrated density into the components of opacity area and magnitude independently. From this figure it can be observed that lens opacity area and magnitude reach asymptotes quickly at ~25 and ~15 minutes respectively, for mice recovered in the open chamber with unprotected eyes.

To better compare the changes observed in the anterior segment by SD-OCT, the first derivative was taken of the fitted data from **Figures 5C-J**. **Figure 6** demonstrates that changes observed in the unprotected eyes of the mice recovered in the open chamber are more prominent than the other three treatment conditions. In terms of magnitude and duration, the anterior chamber depth has the largest and most sustained rate of change over the post-EPIP recovery period. At 80 minutes post, anterior chamber depth changes persist and have yet to reach an asymptotic limit. The sustained changes occurring in the anterior chamber depth persisted longer than the smaller magnitude responses observed with corneal thinning and lens media opacity integrated density that reached asymptotes at ~25-30 minutes, prior to the earliest documented lesion development at 45 minutes.

**Figure 6.**
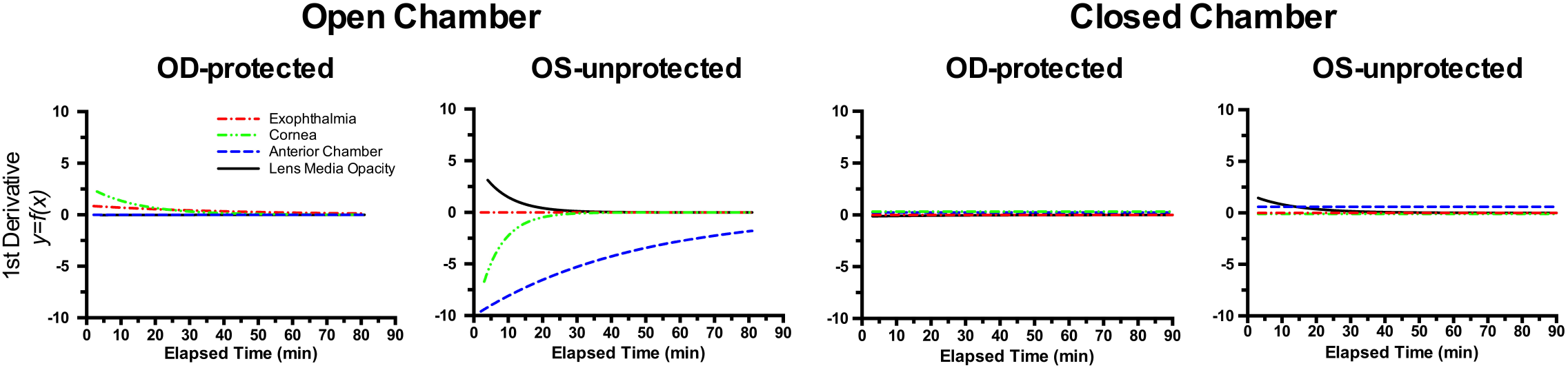
Rate of Change Comparisons. First derivatives taken of the fitted curve trends shown in **Fig. 5** reveal the rate of change for exophthalmia, corneal thinning, anterior chamber depth collapse, and ocular lens media opacity development. These curves show the magnitude, direction and rate of change associated with these metrics measured *in vivo* via anterior segment SD-OCT imaging. Note that the most prevalent changes occur in the unprotected left eyes (**OS-unprotected**) of mice that are recovered in the open chamber. Furthermore, anterior chamber depth is observed having the largest magnitude and most sustained rate of change during the 80 minutes of monitoring.

**Figure 7.**
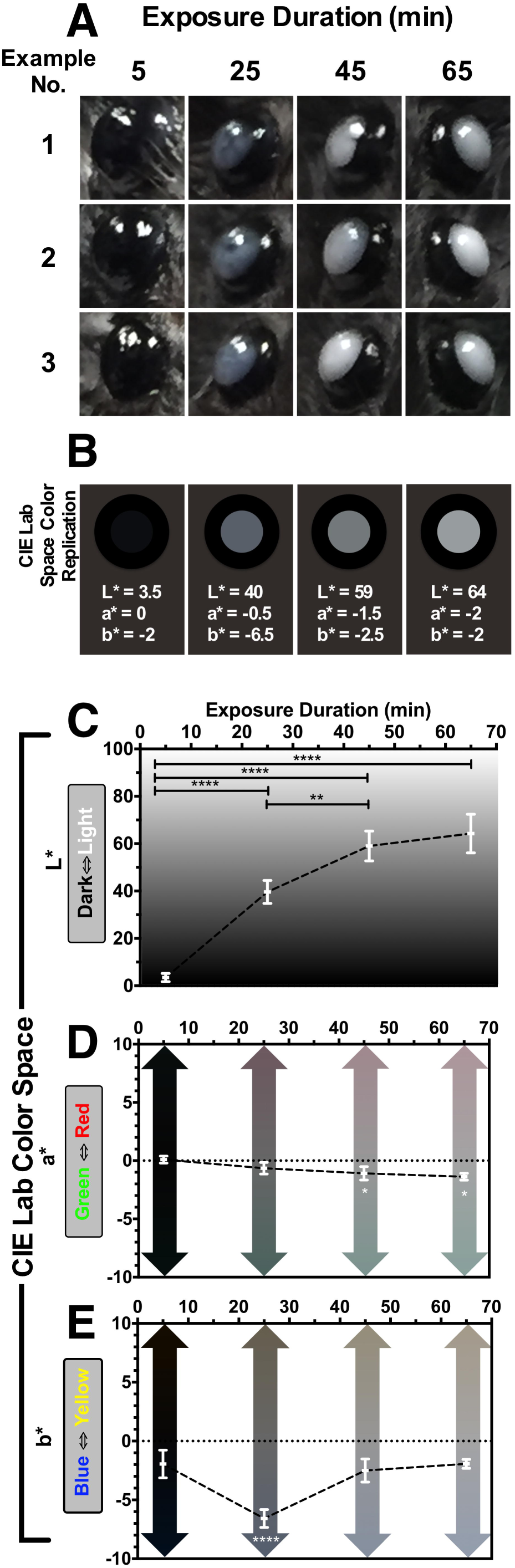
Changes in Ocular Appearance with Exposure Duration. (**A**) Three examples of media opacities from mouse eyes at 5, 25, 45, & 65 minutes post-EPIP. Qualitatively, it is easily observed that eyes immediately following EPIP have a charcoal black appearance and transition to a bluish-gray hue within 25 minutes. Past 25 minutes, eye opacities become brighter in appearance and proceed towards a neutral, grayish-white color. These visual, qualitative observations were quantified so that they could be shown graphically in CIE L*a*b* color space (**B**). (**C**) Brightness (L*) significantly increased with exposure duration. (**D**) Green-Red (a*) exhibited a small (~2-3 units), but significant shift from neutral to a green hue after 45 and 65 minutes. (**E**) Blue-Yellow (b*) experienced a slightly larger (5 units) and very significant shift (p<0.0001) to blue at 25 minutes from the original neutral black color at 5 minutes. Moreover, at 45 & 65 minutes post-EPIP, the blue hue found at 25 minutes significantly returned to the original baseline value observed at 5 minutes. Color values obtained for the plots shown in **C-E** were used to generate a CIE Lab Space Color Rendition **(B)** to artificially recreate the mean appearance of mice with ocular opacities. In these examples the mean CIE Lab values for the pupil are reported as L*, a*, b* values and displayed in a two-dimensional, en face view of the mouse eye including the surrounding iris and periorbital region. This example demonstrates that the reconstituted color values are similar to the in vivo digital color photographic observations and accurately replicate these changes. Retinal lesion impact area correlated moderately strong with ocular exposure duration, decreasing anterior chamber depth and opacity brightness. Thus, it would appear that opacity brightness is a good visual indicator of lesion development probability.

### Visual Recognition of Ocular Lens Position by Visual Assessment of Media Opacity Color and Brightness

**Figure 7** shows color digital photos of the eyes of mice that were enrolled in the interrupted recovery experiments. Three examples per time point show the media opacities at 5, 25, 45, and 65 minutes post-EPIP. **Fig. 7A** demonstrates that the color and brightness of the media opacity changed over time with ocular exposure duration. The quantitative data extracted from the media opacity images is shown in **Fig. 7B-D** and presented in CIE L*a*b* color space. In L* (**Fig. 7B**), the brightness of the media opacity increased significantly with exposure time until reaching the 45 to 65 minute data points where it appears to be approaching an asymptotic limit. In a* (**Fig. 7C**), the neutral green-red level observed at 5 and 25 min post-EPIP changes significantly to a more green hue at 45 (Adj. p=0.0210) and 65 (Adj. p=0.0127) minutes. With b* (**Fig. 7D**), the very slight blue hue observed at 5 min post-EPIP changes significantly more blue at 25 minutes (Adj. p<0.0001), then significantly returns to the original baseline level observed at 5 min for the 45 (Adj. p<0.0001) and 65 (Adj. p<0.0001) minute data points.

To determine whether the mean CIE L*a*b* values obtained accurately represented the digital photos; a color rendition was generated for the four time points evaluated. Shown below the representative mouse eye media opacity photos in **Figure 7A** are the color renditions, which appear to accurately represent what is observed in the digital color photos of exposed mouse eyes.

### Pearson Correlation Results from Interrupted Recovery Experiments

A Pearson correlation test showed that the mean collective lesion impact area values correlated moderately strong with exposure duration (r=0.67; p=0.0018), anterior chamber depth (r=-0.63; p=0.005) and CIE L* brightness (r=0.56; p=0.0452). All other variables measured were insignificant, including exophthalmia, lens opacity magnitude, corneal thickness, and CIE a* & b* trends.

## Discussion

Numerous adverse effects, some of which may render subjects vulnerable to unforeseen complications, have been reported in mice anesthetized with the popular mixture combination of Ketamine and Xylazine. These include hypothermia, bradycardia, hypoxia, and alteration of blood-gas tensions such as acute respiratory acidosis and hypercapnia (Arras, Autenried et al. 2001, Tsukamoto, Serizawa et al. 2015). Retina and brain are two of the most highly metabolic organs in the entire body (Wong-Riley 2010). Surprisingly, when performing experiments on small animals, often little or no proactive measures are used to counter these adverse effects on normal physiology of these tissues.

In terms of vision related complications, practically any form of general anesthesia will also have profound effects on the delicate tissues of the eye. Fragile ocular surfaces exposed to air rely heavily on eyelid function and tear film replenishment for the preservation of corneal integrity (Peng, Cerretani et al. 2013). Cessation of an involuntary blink reflex, which occurs rapidly with the onset of general anesthesia, means tear film depletion and corneal desiccation are imminent. Exophthalmia further exacerbates this problem as increasing palprebal space accelerates the rate at which desiccation occurs (Rolando and Refojo 1983). Although not considered extreme, standard environmental room conditions (~25°C & 20-45% relative humidity) found in practically all climate-controlled laboratories can still be extremely deleterious to the eye if left exposed for a prolonged period. In addition to these visibly apparent side effects, many undesirable ocular changes have been reported in mice following KX anesthesia including reversible cataracts or media opacities (Weinstock and Scott 1967, Bermudez, 2011 #1121, Calderone, Grimes et al. 1986, Ridder, Nusinowitz et al. 2002, Bermudez, Vicente et al. 2011, Bell, Kaul et al. 2014), reduced intraocular pressure (Avila, Carre et al. 2001) (Ding, Wang et al. 2011), corneal damage (Turner and Albassam 2005, Koehn, Meyer et al. 2015), refractive shift (Tkatchenko and Tkatchenko 2010), and compromised retinal and choroidal perfusion (Muir and Duong 2011, Moult, Choi et al. 2017). As others and we (**Fig 1A & B)** have shown, many of these effects are short-lived and usually reverse upon recovery; however, long-term damage can ensue.

In these studies we have demonstrated that lesions involving the outer retina develop in two independent lines of mice following general anesthesia and simulated routine experimental procedures. The two recovery conditions tested demonstrate that: (1) eyes protected from desiccation using evaporation-impermeable methods (ointment + eye shield) do not result in the development of any immediate, or latent, ocular complications regardless of how the mouse is recovered (open vs. closed chambers), and (2) eyes left unprotected, but insulated from the effects of evaporation accomplished by placing the mouse in a closed, humidified chamber showed only one minor complication (lens media opacities) that resolved upon recovery. This in direct contrast to eyes left unprotected and exposed to the effects of evaporation, accomplished by placing the mouse in the open recovery chamber, which exhibited a multitude of ocular changes and complications including prominent retinal lesions. Eyes left exposed were subjected to prolonged desiccative effects of circulating room air at standard environmental temperature and humidity levels. A sigmoid curve fit of the data obtained in this study shows that the risk initiates around 30 minutes post-exposure onset and reaches a maximum probability of ~90% at ~45 minutes.

The comprehensive imaging studies we performed involving both anterior and posterior poles provided probable cause as to why a lesion did or did not materialize under the contrasting conditions tested. As anterior segment SD-OCT imaging showed, mice that recovered in the open chamber, without any ocular protection, underwent continual change over time as the sedation cycle ran its natural course. Our original hypothesis (lesions were caused by exophthalmia) was negated by observations of ocular exophthalmia in all mice regardless of ocular protection or recovery chamber status (**Fig. 5C & 5D)**. Proptosis occurred soon after recumbence and essentially remained constant, relative to other parameters measured, over the time period the animals were followed post-EPIP (**Fig. 6**). After anesthesia induction, visible exophthalmia was observed and soon followed concomitantly by other changes such corneal thinning, lens media opacity changes, and anterior chamber depth. Lens media opacity magnitude appears to reach a plateau first at ~15 minutes, followed by corneal thinning at ~20 minutes and lens media opacity area and integrated density at ~30 minutes. These changes all reached asymptotes at 15-30 minutes post-recumbence with exception to anterior chamber depth, which continued to change with increasing exposure time. This decrease did not seem to occur as a result of the cornea collapsing and/or the deflation of anterior chamber compartment. Instead, the lens steadily migrated into the anterior chamber as a result of a void being left by the loss of aqueous humor. Previous studies with KX anesthesia in mice have shown a precipitous decline in intraocular pressure over time (Ding, Wang et al. 2011), which tends to support our imaging observations of decreasing anterior chamber depth that can be extrapolated as being synonymous with decreasing anterior chamber volume. As revealed by **Figures 5A & 5G** lens migration was substantial and approximately half of the lenses observed came into direct contact with the posterior corneal surface. In some occurrences, the anterior lens capsule appeared to adhere to the cornea endothelium (**Suppl. Fig. S6-Ex. A**-arrows) causing traction on the lens capsule and opening a void filled with semi-reflective fluid between the capsule and lens (**Suppl. Fig. S6-Ex. A**-asterisks). At long exposure times (~1 hr or more), any aqueous humor remaining within the anterior chamber became semi-reflective by anterior segment SD-OCT; presumably due to precipitated analytes, cellular infiltration, or protein flare (**Suppl. Fig. S6-Ex. B-asterisks**). Concomitantly during this time, it could be observed that perturbations in shape and symmetry of the cornea began to emerge (**Suppl. Fig. S6-Ex. A&B)**. We suspect that these anterior segment changes are substantial enough that they could be the underlying cause of corneal ulcerations and microphthalmia that commonly occur in mice following experimental studies; which occurred in 14% (3/22) of our mice left exposed in the open chamber for 65 minutes or longer.

Based on these observations we propose that a cascade of events leads to the formation of retinal lesions in unprotected eyes. First, when mice are administered anesthesia and pupil dilation drops, they receive an extremely large dose of alpha-1 & 2 adrenergic receptor agonists that results in vasoconstriction, extraocular muscle relaxation, and as we and others have shown, exophthalmia (Calderone, Grimes et al. 1986). Soon thereafter, processes involved with tear film production and aqueous fluid turnover are suspected to be “clamped” or at least substantially disrupted (Calderone, Grimes et al. 1986). These conditions render the eye prone to complications, as it is unable to regulate and properly supply and/or drain aqueous humor production from the ciliary bodies, Trabecular meshwork and Schlemm’s canal. Drug-induced exophthalmia, causing excessive ocular exposure, results in rapid depletion of the tear film whereby corneal desiccation and dehydration ensue. As the rate of evaporation exceeds that of aqueous humor production, prolonged exposure depletes aqueous humor volume via transcorneal water loss (Weinstock and Scott 1967, Fraunfelder and Burns 1970). As transcorneal water loss progresses, the steady reduction in anterior chamber volume is subsequently followed by a concomitant decrease in intraocular pressure. As aqueous volume and pressure decline, we speculate that a pressure imbalance occurs between the anterior and posterior chambers. This causes the lens to be either drawn into the anterior chamber by a negative pressure created by the depletion of aqueous humor or alternatively, the lens may be pushed into the anterior chamber by the positive pressure that remains within the posterior chamber. Alternatively, exophthalmia appears to position the lens equator right at the supraorbital margins. Taking into account the decreasing anterior chamber volume and pressure, in conjunction with the marble-like rigidity of the lens, this atypical positioning arrangement of the globe may apply enough extra-orbital pressure on the superior and inferior regions to semi-extrude or propel the lens into the anterior chamber.

As exposure duration progresses, the prolapsing lens begins to apply ever-increasing traction on the radial suspensory ligaments (e.g. zonules of Zinn). The ligaments connect the lens to the inner ocular surface, inserting first at the ciliary processes, passing through the Pars plana, and terminating at the Ora serrata (McCulloch 1954, Shi, Tu et al. 2013). The ligaments tether the lens to the inner ocular globe surface in a completely circumferential manner. As the lens moves farther into the anterior chamber, the force being applied on RPE-Bruch’s membrane complex exceeds the point at which some areas of the choriocapillaris can sustain perfusion, ultimately, causing localized ischemia to certain regions. The ischemia is short-lived as it is followed immediately thereafter by reperfusion upon recovery from the effects of sedation. Unfortunately, the cause and effect outlined here remains speculative since it is impossible to visualize this process in real time by *in vivo* imaging. Non-invasive imaging methods (e.g. FA, ICGA or OCT-A) that could be used to further investigate this phenomenon are not feasible due to the severe media opacities that occur during the process.

Imaging and histology observations have revealed that the induced damage is limited to the outer retina as no evidence of retinal vasculature or inner retina damage has been documented. The damage we have observed post-induction via non-invasive imaging appears to be specifically limited to the RPE and immediately adjacent photoreceptor layer. This observation underscores how important perfusion and oxygenation are to the metabolically demanding outer retina and RPE. The diminishing lesion visibility over time suggests that there is an initial area at risk followed by a smaller area of necrosis, much like that observed in ischemia-reperfusion injury of mouse myocardial tissue (Bohl, Medway et al. 2009). In the long-term example shown in **Figure 4**, the dark area demarcated at day 3 by IRDF was suggestive of an area at risk as the visible IRDF changes disappeared by 14 days post-induction. Meanwhile, RFDF, BAF, TEFI color fundus, and FA-SLO show that perturbations still persist at 80 days post-induction and are indicative of some lasting consequences. Between these time points the initial area of risk became an area of necrosis that was only ~75% of the original risk size. Ideally, future studies would address in more detail the dynamics between acutely visible damage and the long-term consequences of the initial brief insult observed at 3 days post-induction.

Finally, we observed an interesting trend in the color/brightness of media opacities and the risk of lesion formation. If the opacity is of faint brightness, and has a bluish-gray hue, then our observations demonstrate that there is a low risk that a retinal lesion has formed by that particular moment. However, if the media opacity is bright, and predominantly more grayish-white in hue, then it is very likely that the retina is at risk of developing lesions. These changes appear to occur in two general phases that can be divided at ~25-minute recovery mark. Under ½ hour, during the early phase, opacity magnitude and area reach a plateau within ~25 minutes. At this time the reduction (~25%) in anterior chamber depth is small and thus the lens is still quite distal from the posterior cornea. This condition gives the lens media opacity a bluish hue to the observer since the lens is still quite distal from the cornea. As we showed in **Fig. 5** & 6, beyond ½ hour during the late phase the media opacity magnitude and area are no longer changing since they have reached plateaus. Still changing in the late phase however, is the anterior chamber depth and lens position, which has now further halved the distance to the posterior cornea with 20 additional minutes elapsed by the 45 min time point. This change now places the lens within 150 μm (range ~75-200 μm) of the posterior cornea, thus causing the media opacity to lose its bluish-hue and become brighter and grayish-white in appearance. This trend continues beyond 45 minutes with opacities becoming brighter and slightly whiter as the lens continues to approach the cornea. In summary, mice with eyes that look like white ping-pong balls have an extremely very high probability of exhibiting retinal lesions.

In this study we provided strong evidence to show that spontaneous retinal lesions can occur in mice simply from undergoing anesthesia induction and experimental manipulation followed by inadequate post-procedure care. These observations were supported by substantial structural and functional evidence. Lesion severity diminishes rapidly within about 2 weeks but evidence of long-term damage persists at 2.5 months post-induction. This discovery is highlighted by the fact that these are previously undocumented observations that could potentially be useful to researchers wishing to induce transient ischemia-reperfusion injury to the outer retina of small rodents. Since the currently observable damage appears to be limited to the RPE and outer retina it is plausible that this model is unique from previously published methods used achieve retinal ischemia in mice by elevating intraocular pressure or performing arterial ligation (Buchi, Suivaizdis et al. 1991, Minhas, Morishita et al. 2015, Hartsock, Cho et al. 2016). As we have shown, this model is easily reproduced and is unique from previous approaches as it can be non-surgically and non-invasively induced. In closing, additional studies are warranted to determine the full ramifications of the long-term damage on the retina following these acutely induced lesions.

## Acknowledgements

We thank Gayle Pauer, Charlie Kaul, Rupesh Singh, Matt Ford, and Ibraham Seven for technical assistance and constructive comments.

## Supplementary Figures

**Figure S1 – Exophthalmia Induced by Adrenergic Agonists Phenylephrine and Xylazine**

**Figure S2 – Uninterrupted Recovery Experiment Results**

**Figure S3 – ERG Results at 30-days Post-EPIP**

**Figure S4 – TEFI Color Fundus Lesion Examples at 3-days Post-EPIP**

**Figure S5 – Lens Media Opacity Area and Magnitude**

**Figure S6 – Additional Anterior Segment Observations made with SD-OCT: Misshapen Cornea, Semi-reflective Media in the Anterior Chamber and Under the Lens Capsule Concomitant with Lens Capsule Adhesions to the Posterior Cornea.**

