## Supplementary figures and images for "Prolonged Ocular Exposure Leads to the Formation of Retinal Lesions in Mice"

### Supplemental Figure 1

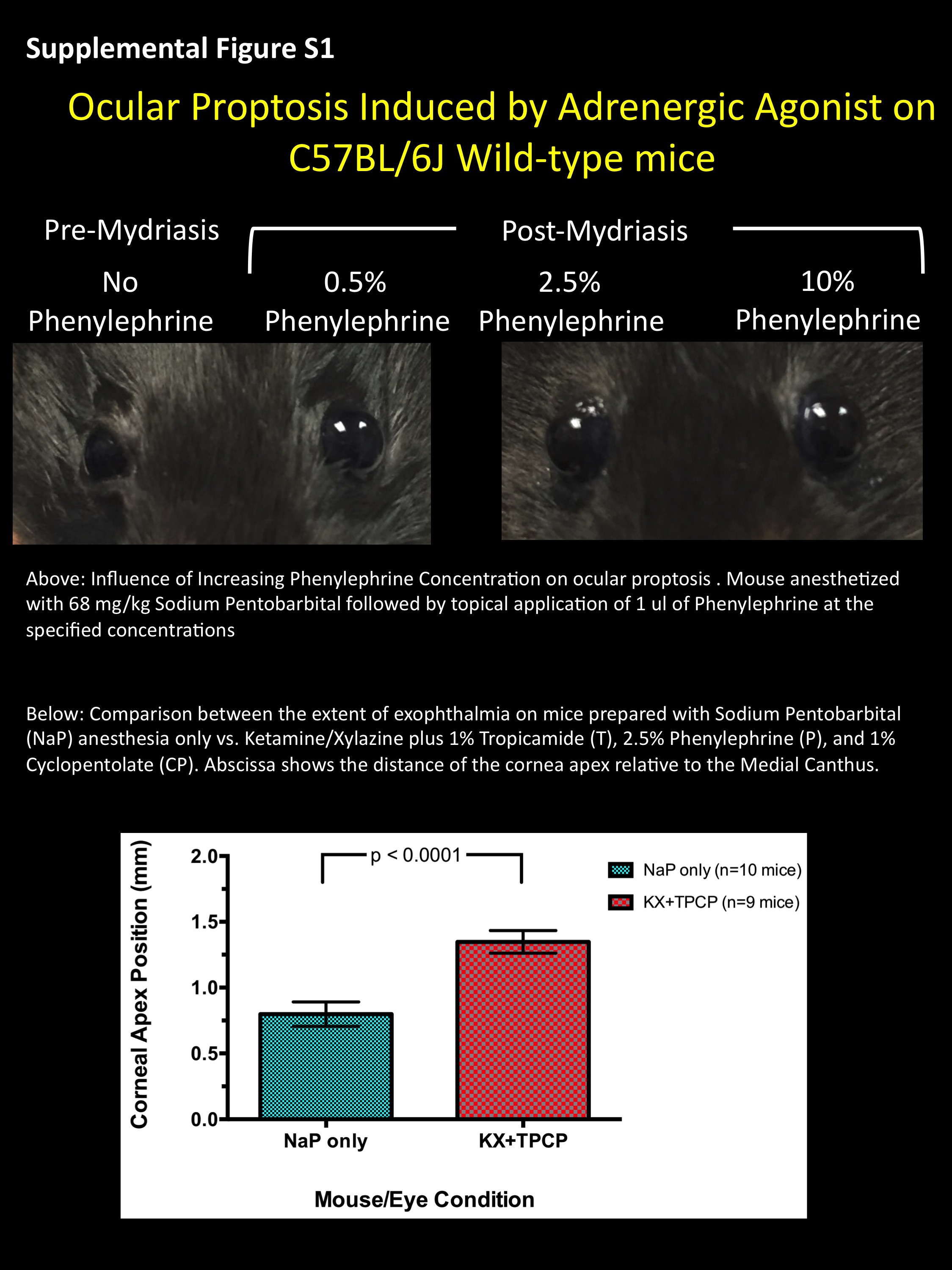

### Supplemental Figure 2

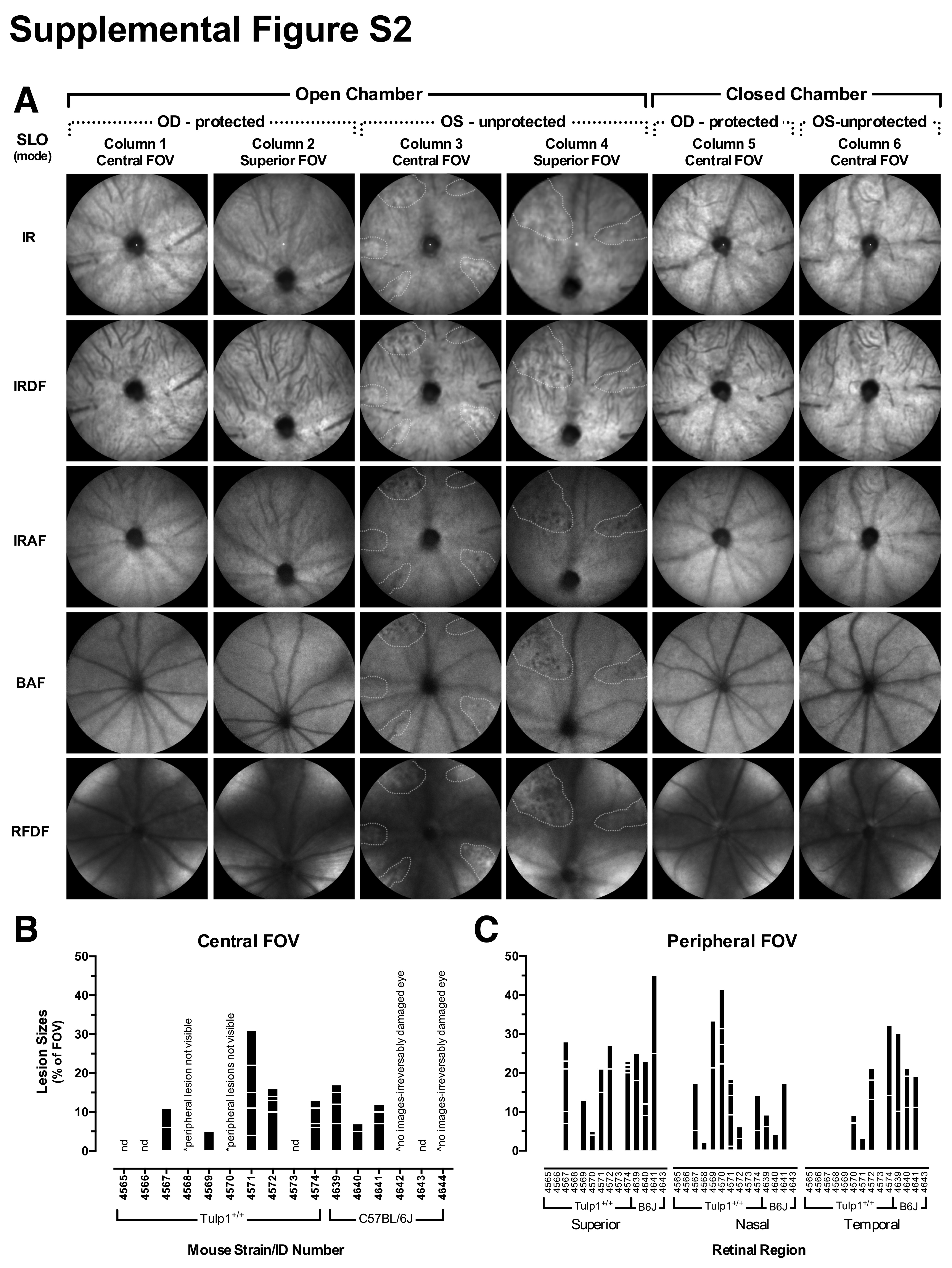

### Supplemental Figure 3

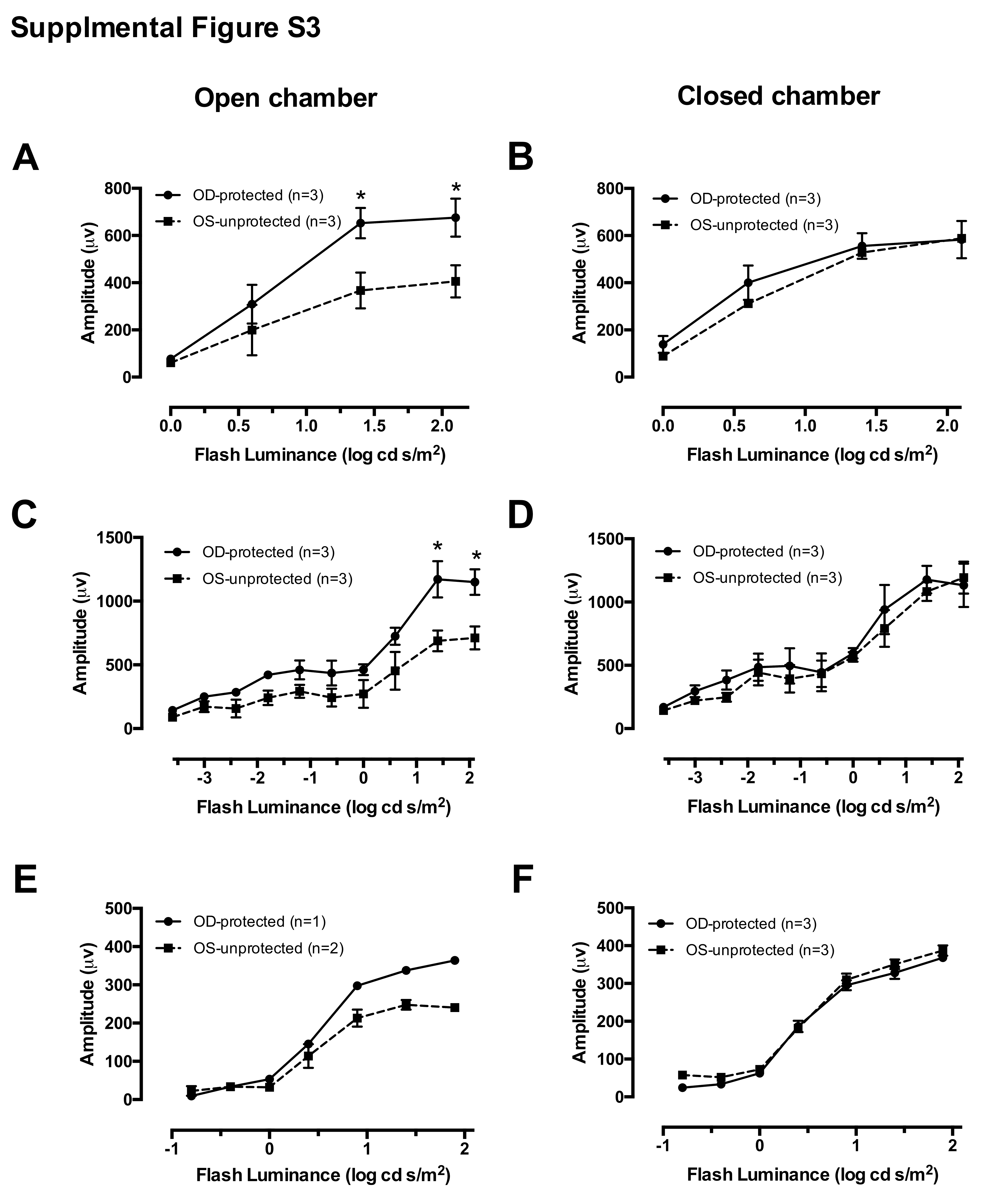

### Supplemental Figure 4

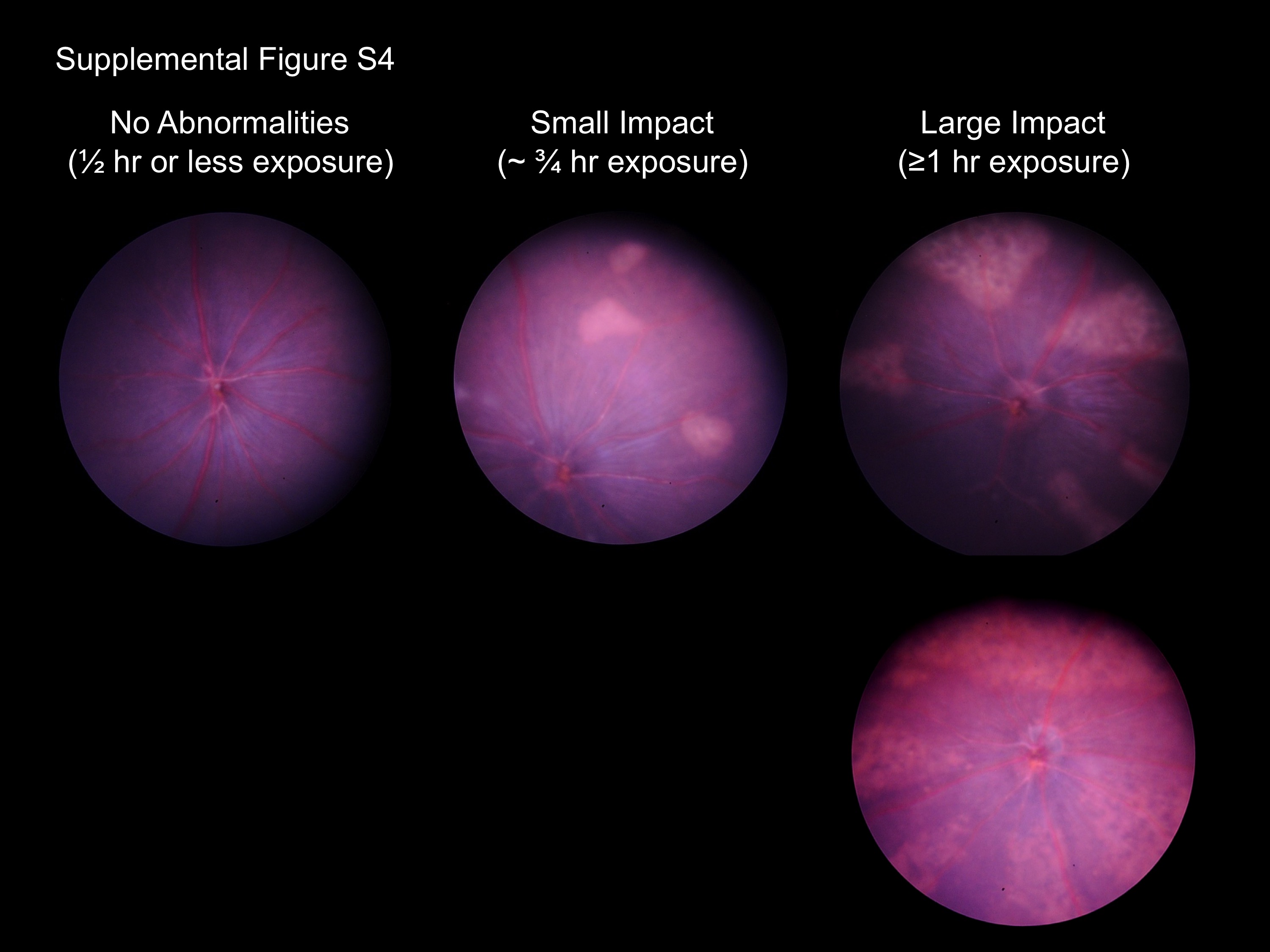

### Supplemental Figure 5

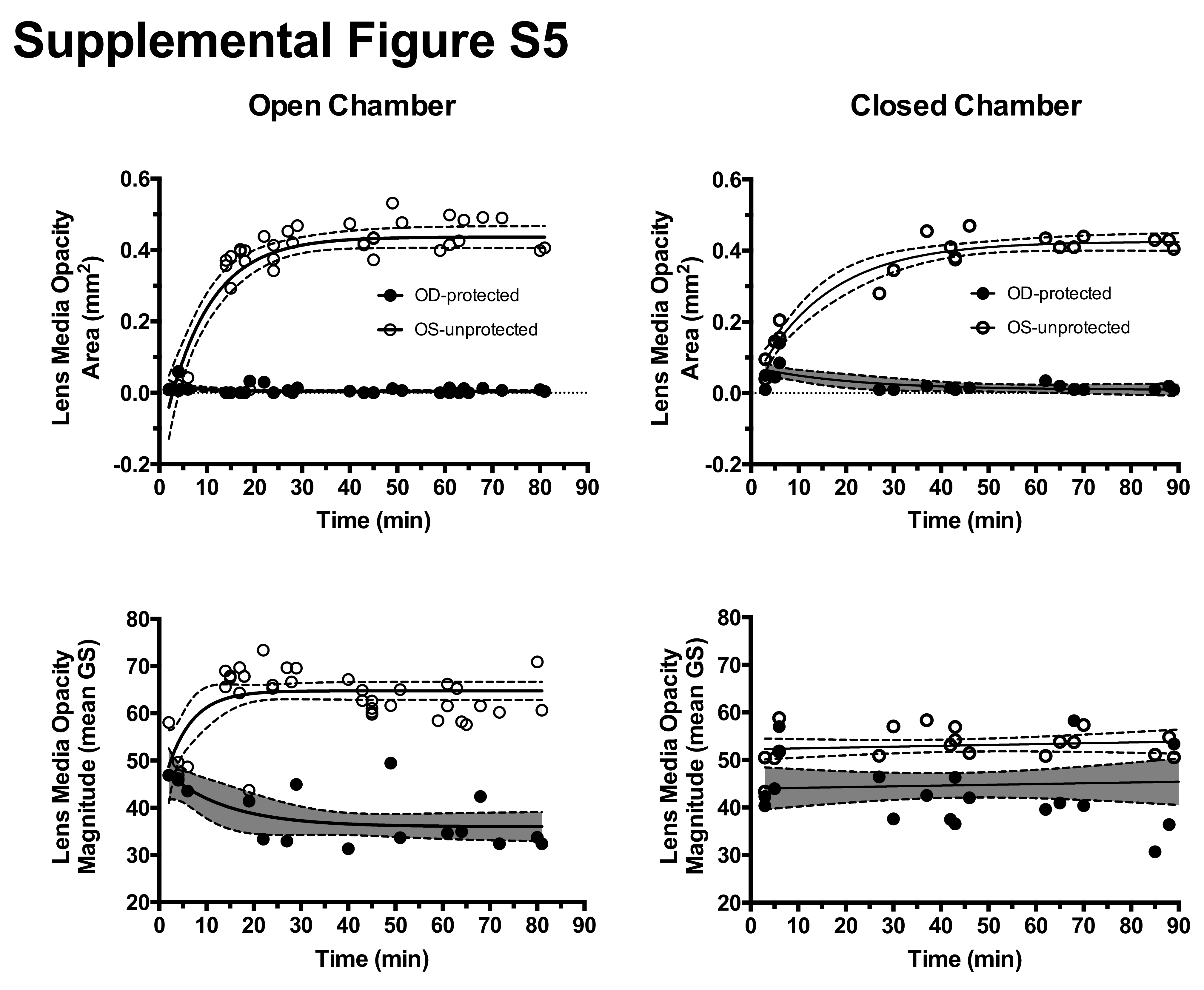

### Supplemental Figure 6

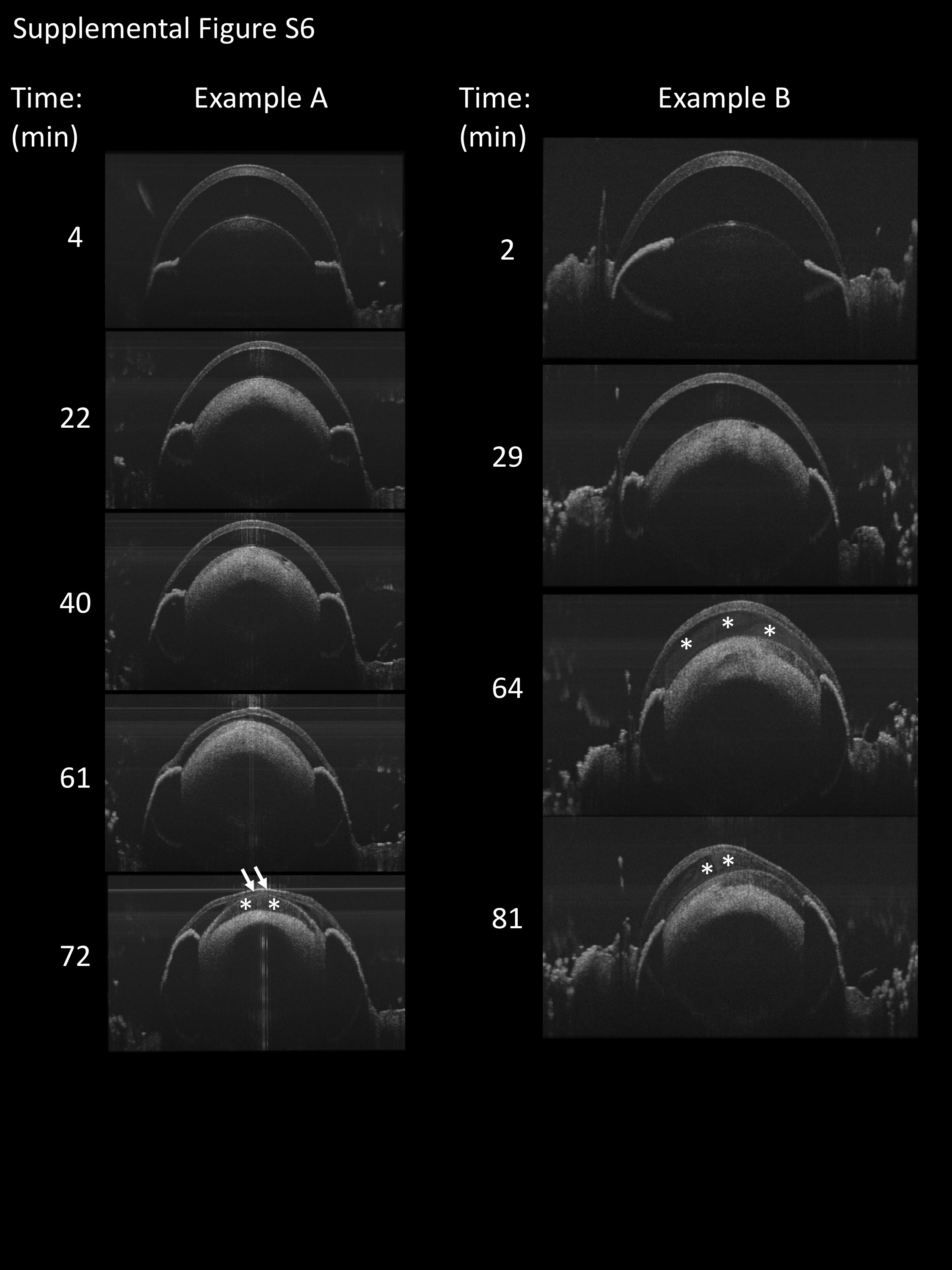
